# Several phased siRNA annotation methods can frequently misidentify 24 nucleotide siRNA-dominated *PHAS* loci

**DOI:** 10.1101/409417

**Authors:** Seth Polydore, Alice Lunardon, Michael J. Axtell

## Abstract

Small RNAs regulate key physiological functions in land plants. Small RNAs can be divided into two categories: microRNAs (miRNAs) and short interfering RNAs (siRNAs); siRNAs are further sub-divided into transposon/repetitive region-localized heterochromatic siRNAs and phased siRNAs (phasiRNAs). PhasiRNAs are produced from the miRNA-mediated cleavage of a Pol II RNA transcript; the miRNA cleavage site provides a defined starting point from which phasiRNAs are produced in a distinctly phased pattern. 21-22 nucleotide (nt)-dominated phasiRNA-producing loci (*PHAS*) are well represented in all land plants to date. In contrast, 24 nt-dominated *PHAS* loci are known to be encoded only in monocots and are generally restricted to male reproductive tissues. Currently, only one miRNA (miR2275) is known to trigger the production of these 24 nt-dominated *PHAS* loci. In this study, we use stringent methodologies in order to examine whether or not 24 nt-dominated *PHAS* loci also exist in *Arabidopsis thaliana*. We find that highly expressed heterochromatic siRNAs were consistently mis-identified as 24 nt-dominated *PHAS* loci using multiple *PHAS*-detecting algorithms. We also find that *MIR2275* is not found in *A. thaliana*, and it seems to have been lost in the last common ancestor of Brassicales. Altogether, our research highlights the potential issues with widely used *PHAS*-detecting algorithms which may lead to false positives when trying to annotate new *PHAS*, especially 24 nt-dominated loci.

## Introduction

Small RNAs regulate key physiological functions in land plants, ranging from organogenesis (Boualem et al., 2008, 2008; Kutter et al., 2007; Laufs et al., 2004; Williams et al., 2005) to gametogenesis (Grant-Downton et al., 2009). Three major protein families are involved in the biogenesis of small RNAs. The first family is the DICER-LIKE (DCL) protein family. Consisting of four paralogs in the *Arabidopsis thaliana* genome (Baulcombe, 2004; Chapman & Carrington, 2007), DCL proteins hydrolyze RNA precursors into 20-24 nt double stranded RNA fragments (Millar & Waterhouse, 2005). These double-stranded RNA fragments are then loaded into ARGONAUTE (AGO) proteins, the second protein family, and one strand of the RNA is discarded (Ender & Meister, 2010). Upon Watson-crick binding to other RNA transcripts in the cell, the AGO/single-stranded RNA complex represses other RNA transcripts (Baumberger & Baulcombe, 2005; Qi et al., 2005). Overall, 10 AGOs are encoded in the *A. thaliana* genome (Tolia & Joshua-Tor, 2007). RNA DEPENDENT RNA POLYMEREASES (RDRs) are the third family of proteins involved in the biogenesis of many small RNAs. RDRs convert single-stranded RNAs into double-stranded RNAs by synthesizing the complementary strand of the RNA molecule (Willmann, et al., 2011). Six *RDRs* are encoded in the *A. thaliana* genome (Willmann, et al., 2011).

Small RNAs can be divided into two major categories: microRNAs (miRNAs) which are precisely processed from single-stranded RNA with a hairpin-like secondary structure (Millar & Waterhouse, 2005), and short interfering RNAs (siRNAs), which are derived from double-stranded RNA precursors (Axtell, 2013). siRNAs are further divided into several different groups, including phased siRNAs (phasiRNAs) and heterochromatic siRNAs (hc-siRNAs). Predominantly 24 nts in length, hc-siRNAs function to repress transcription of deleterious genomic elements such as transposable elements or repetitive elements (Ahmed et al., 2011) and the promoters of certain genes (Baev et al., 2010) by reinforcing the presence of heterochromatin in targeted areas (Baulcombe, 2004; Sugiyama et al., 2005). Biogenesis of hc-siRNAs begins with transcription by the plant-specific, holo-enzyme DNA DEPENDENT RNA POLYMERASE IV (Pol IV) (Onodera et al., 2005). The resulting transcript is then converted into double-stranded RNA by RDR2 and this double-stranded transcript is hydrolyzed by DCL3 (Matzke et al., 2009). phasiRNAs are derived from DNA DEPENDENT RNA POLYMERASE II (Pol II) transcripts that have been targeted by miRNAs (Fei et al., 2013). Upon miRNA-mediated hydrolysis, the RNA transcript is converted into double-stranded RNA by RDR6 (Cuperus et al., 2010). The resulting double-stranded RNA is then cleaved into 21nt double-stranded RNA fragments by DCL4 (and less frequently DCL2) (Axtell et al., 2006).

21-22 nt phasiRNA-producing loci (*PHAS*) are clearly represented in all land plants that have been sequenced thus far (Fei et al., 2013; Zheng et al., 2015). However, 24 nt dominated *PHAS* loci are only currently described in rice (Song et al., 2011), maize (Zhai et al., 2015), and other non-grass monocots (Kakrana et al., 2018). Much like 21 nt-dominated *PHAS*, the biogenesis of these 24 nt-dominated *PHAS* loci begins with the Pol II-dependent transcription of a single-stranded RNA precursor which is then targeted by miR2275 and hydrolyzed. To date, miR2275 is the only miRNA known to trigger the production of 24 nt-dominated phasiRNAs (Fei et al., 2013). The resulting RNA transcript is then converted into a double-stranded RNA molecule by RDR6 (Zhai et al., 2015). However, these phasiRNA precursors are then hydrolyzed by DCL5 (a DCL3 homolog sometimes called DCL3b) to produce 24 nt phasiRNAs (Fei et al., 2013).

Aside from the combination of their size and biogenesis patterns, 24 nt-dominated *PHAS* loci are distinct in various ways. These loci as well as their triggering miRNA, miR2275, are very specifically expressed in the tapetum during early meiosis and quickly recede in expression in other stages of male gametogenesis in rice and maize (Tamim et al., 2018). The AGO protein that loads these phasiRNAs is unknown; however, in maize, *AGO18b* expression levels match those of the 24 nt-dominated *PHAS* loci quite closely and is therefore the most likely candidate to load 24 nt phasiRNAs (Komiya et al., 2014; Zhang et al., 2015). The targets of these 24 nt phasiRNAs are unknown, but they are apparently necessary for proper male gametogenesis (Ono et al., 2018). 24 nt-dominated *PHAS* loci were also described in the non-grass monocots asparagus, lily, and daylily (Kakrana et al., 2018). These phasiRNAs are produced from processing of inverted repeat (IR) RNAs, instead of the double-stranded RNA precursors observed in rice and maize (Kakrana et al., 2018). Although the 24 nt-dominated *PHAS* loci from non-grass monocots are still expressed most greatly in male reproductive tissue, in asparagus they are also expressed in female reproductive tissue (Kakrana et al., 2018).

We set out to search for evidence of 24 nt-*PHAS* loci in plants besides monocots. We searched for 24 nt *PHAS* loci in the *A. thaliana* genome using small RNA-seq data. Currently, several distinct algorithms are available to calculate the “phasing” of a sRNA-producing locus (Dotto et al., 2014; Guo et al., 2015; Zheng et al., 2014). In general, these algorithms calculate the number of reads that are “in-phase” against those that are “out-of-phase” in order to determine the likelihood that a particular locus truly produces phasiRNAs (Axtell, 2010). However, 24 nt-dominated siRNA loci are very numerous in angiosperms, and therefore are a potential source of false-positives during searches for *PHAS* loci. We therefore carefully examined *A. thaliana* 24 nt-dominated loci that consistently passed *PHAS*-locus detecting algorithms using multiple methods find that they are likely just heterochromatic siRNAs (hc-siRNAs). We also use two other methods to examine the presence of 24 nt-dominated *PHAS* loci in the *A. thaliana* genome. We searched for *rdr6*-dependent, 24 nt-dominated loci and found 18 such loci. We also examined homology of the miR2275 which triggers 24 nt phasiRNA biogenesis in rice and maize but found that the Brassicales clade contains no potential homologs for this miRNA. Overall, our results suggest that there are no true 24 nt *PHAS* loci in *A. thaliana*. Furthermore, our analysis shows that existing phasing score algorithms to detect novel *PHAS* loci can lead to false positives.

## Results

### miR2275 is not found in the *Brassicales* clade

Currently known 24 nt dominated phasiRNA precursors are known to be targeted only by a single miRNA family, miR2275 (Song et al., 2008; Zhai et al., 2015). We examined all available angiosperm genomes on Phytozome (ver 12.1) for potential homologs of *MIR2275*. In monocots, all but *Brachypodium stacei, Spirodela polyrhiza* and *Zostera marina* had potential miR2275 homologs (Figure 1; Figure S1). 12 eudicots had potential miR2275 homologs based on sequence similarity (Figure 1). We therefore interrogated small RNA libraries for these species to see if a mature miR2275 homolog was expressed. Five eudicots had evidence of mature miR2275 accumulation (Figure 1). All of the monocots for which we obtained small RNA-seq data expressed mature miR2275, except *Musa acuminata* (Figure 1). Alignment of the *MIR2275* loci in species for which potential homologs could be identified shows strong conservation of the mature miRNA and miRNA* sequences (Figure S2), suggesting that these loci evolved from a common ancestor. Importantly, because of the high specificity of miR2275 expression in developing anthers (Tamim et al., 2018), it’s possible that our analysis includes false negatives, especially in situations where no reproductive tissue small RNA libraries were available. Altogether, our data suggest that miR2275 is not found in *A. thaliana* and that this loss apparently occurred before the last common ancestor for Brassicales.

**Figure 1.**
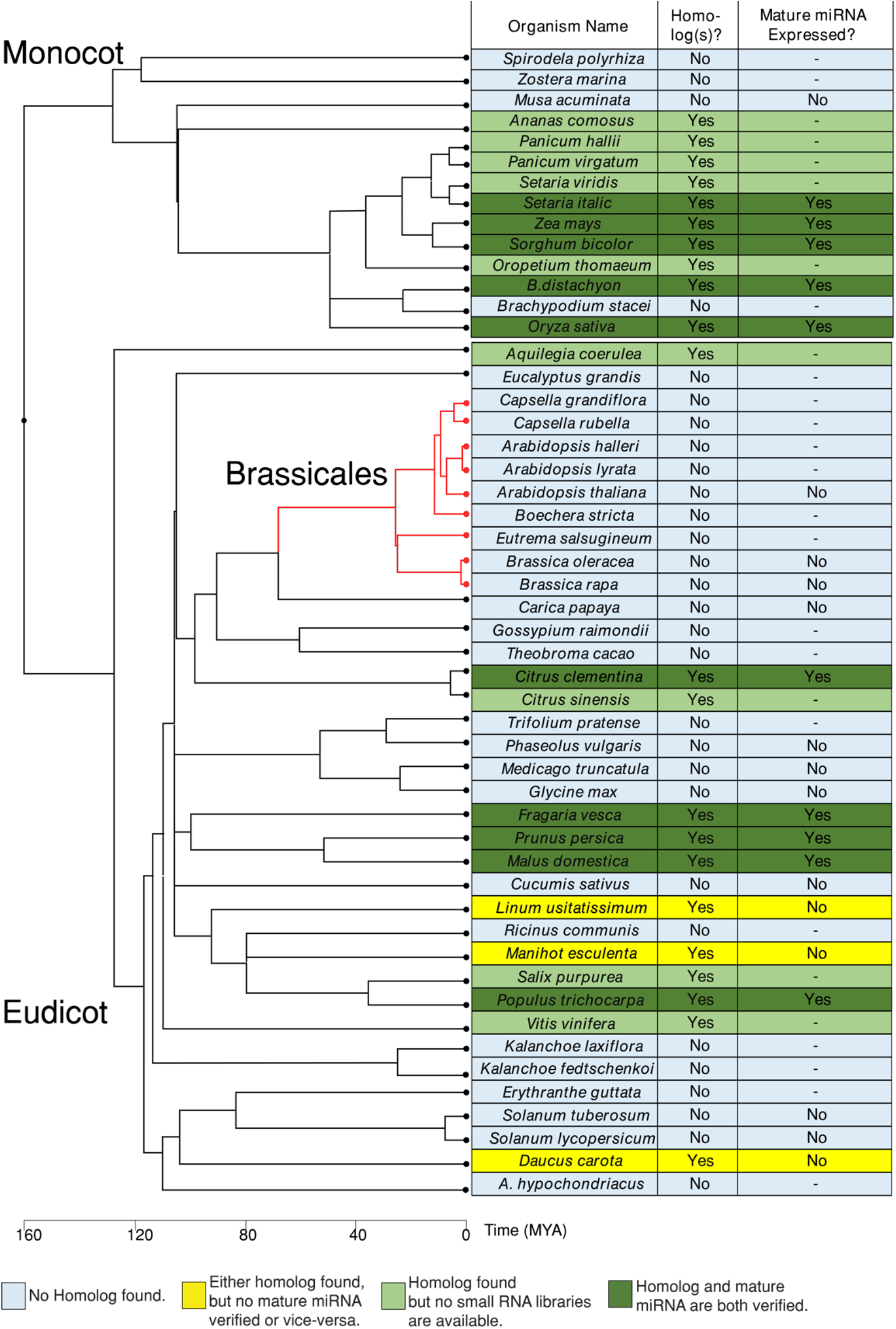
Conservation of *MIR2275*. Evidence for existence of *MIR2275* homologs in angiosperms. Phylogeny depicts estimated divergence times per TimeTree of Life (Kumar et al., 2017).

### Three *A. thaliana* hc-siRNA loci consistently pass *PHAS*-detecting algorithms

Although *A. thaliana* lacks miR2275, it’s possible that 24 nt *PHAS* loci exist in *A. thaliana* and are triggered by a different small RNA. We reasoned that true 24 nt *PHAS* loci would be dependent on one or more of the well-described *A. thaliana RDR* genes: *RDR1, RDR2*, or *RDR6*. We thus examined a previously described differential expression analysis that identified *A. thaliana* small RNA loci that were down-regulated in an *rdr1/rdr2/rdr6* triple mutant (Polydore & Axtell, 2018). The phase scores of *rdr1/rdr2/rdr6*-dependent, 24 nt-dominated loci were calculated in eight independent small RNA libraries using three different algorithms (Figure 2a). Reasonable cut-offs for *PHAS* loci detection were determined by examining the phase score distributions in the three merged wild-type libraries when well-known 21 nt *PHAS* loci were analyzed (Figure S3). These cutoffs were consistent when a larger number of sRNA-seq libraries were examined (Figure S4). Of the 31,750 loci examined, only three (Figure S5; Dataset S1) passed the *PHAS* loci algorithms consistently in all 8 libraries examined (Figure 2a).

We were interested in determining why these three loci consistently pass the *PHAS*-detection algorithms. As phasiRNA precursors are known to be targeted by miRNAs, we predicted whether or not any known *A. thaliana* miRNAs could target these three loci. We were unable to find any obvious miRNA target sites at these loci. Although miRNA target sites were not apparent at these loci, it’s entirely possible that other siRNAs might target them and initiate siRNA phasing. We therefore attempted to determine the most common phase register at each of these loci in the 8 different wild-type libraries (Figure S6). If these loci are truly phased, then we would expect the same phase register to predominate in each library examined. While this was the case for *TAS2* (a positive control), none of the three 24 nt loci passing the *PHAS*-detection algorithms had consistent phase registers (Figure S6). We also note that these three loci didn’t have the majority of their mapped reads falling into a single phase register in any of the libraries examined, as one would expect for a true *PHAS* locus (Figure S6).

**Figure 2.**
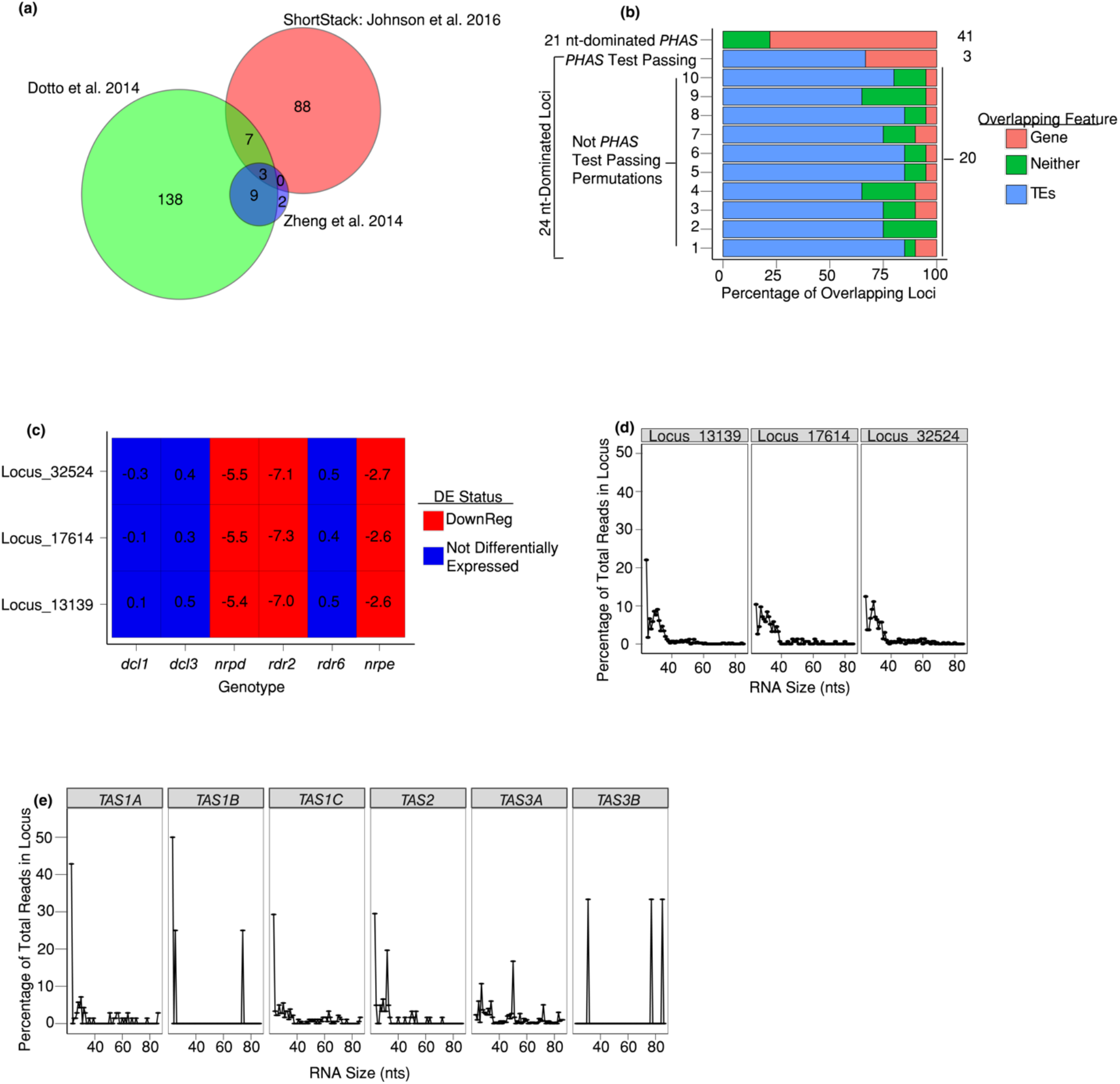
Properties of three *A. thaliana* 24 nt-dominated small RNA loci that were called ’phased’ by three different methods. (a) Venn diagram shows numbers of 24 nt-dominated loci that were called ‘phased’ by the indicated algorithms. (b) Percentage of the three *PHAS*-test passing loci overlapping either genes or transposable elements. The percentage is calculated as: (number of loci intersecting a feature/total number of loci in category)*100. The total number of loci in each category is given on the right. For 24 nt not-*PHAS* loci, ten randomly selected cohorts of 20 loci each are shown. (c) Accumulation of the three *PHAS*-test passing loci in different genetic backgrounds. Numbers represent the ratio of small RNA accumulation in the indicated genotypes over that in corresponding wild-type library as computed by DESeq2. The differential expression status was determined via DESeq2 at an FDR of 0.1. (d) Percentage of short RNAs from *dcl2/dcl3/dcl4* triple mutant libraries by read length for the three *PHAS*-test passing loci. (e) Same as in Panel d, except for five known *A. thaliana TAS* loci.

We then determined where in the genome these loci were encoded. Similar to hc-siRNA loci, these three loci primarily overlap with repeat- and transposable-elements (Figure 2b). In comparison, many known *A. thaliana PHAS* loci are primarily derived from genes (Figure 2b). Rice and maize annotated 24 nt *PHAS* loci are derived from long non-coding RNAs that are encoded in regions of the genome that are devoid of protein-coding genes, transposable elements, or repeats (Song et al., 2008; Zhai et al., 2015). We also note that sRNA accumulation from these three loci is down-regulated in *nrpd1-3* (NRPD is the largest sub-unit of Pol IV), *nrpe* (NRPE is the largest subunit of Pol V), and *rdr2* (Figure 2c). hc-siRNAs have the same genetic requirements (Matzke et al., 2009), which strongly suggests that these three loci produce hc-siRNAs. Notably, these loci are not down-regulated in *dcl3* backgrounds. This is in line with past data that show that DCL4 and DCL2 can partially complement production of hc-siRNAs in *dcl3* backgrounds (Gasciolli et al., 2005).

The Pol IV transcripts from which hc-siRNAs are derived are around 26-50 nts in length, and accumulate to detectable levels in the *dcl2-1/3-1/4-2t* (*dcl234*) triple mutant (Ye et al., 2016; Zhai et al., 2015). We therefore analyzed the lengths of reads mapping to our three putative 24 nt *PHAS* loci in *dcl234* triple mutants. We found most reads were less than 40 nts long, indicating that these putative 24 nt *PHAS* loci are associated with short precursors similar to hc-siRNA loci (Figure 2d). In comparison, reads mapping to the known *TAS* loci had a wider range of sizes, ranging from 24-80 nts (Figure 2e). Overall, the size profile of the precursor RNAs further delineate these three loci from known *PHAS* loci.

Finally, we examined the AGO-enrichments of small RNAs from these loci. Canonical hc-siRNAs are loaded into AGO4 in order to repress other loci transcriptionally (Mi et al., 2008). While sRNAs from our three putative 24 nt *PHAS* loci are not particularly enriched in AGO4-immunoprecipitation libraries, known 21 nt *PHAS* loci are quite depleted in the same dataset (Figure S7). We also note that sRNAs from the three putative 24 nt *PHAS* loci are depleted in AGO1 immunoprecipitation libraries (Figure S7), probably owing to the lack of 21 nt sRNAs, which AGO1 primarily loads, produced at these loci (Mi et al., 2008).

### The three *PHAS*-Test passing loci have distinct characteristics

hc-siRNAs are known to produce small RNAs in a very imprecise manner, unlike the largely precise processing of phasiRNAs (Axtell, 2013). We were interested in how three hc-siRNAs could consistently pass three different *PHAS*-detecting algorithms. One simple explanation is possible erroneous placement of ambiguously mapped reads. These multi-mapping reads could cause the *PHAS*-detecting algorithms to overestimate the number of “in phase” reads. We therefore determined the proportion of multi-mapping reads in each locus, but these loci have low proportion of multi-mapping reads compared to other 24 nt-dominated loci (Figure 3a). We note that these three loci are more highly expressed than most other 24 nt-dominated loci and even known *PHAS* loci in *A. thaliana* (Figure 3b). These three loci accumulate to nearly 1,000 RPM, while most other 24 nt-dominated loci and known *PHAS* loci only produce around 1 RPM (Figure 3b). Another interesting feature of these loci is their length. All three of these loci are around 10,000 nts in length, far greater than the 200-800 nt length of canonical *PHAS* loci and most other 24 nt-dominated loci (Figure 3c). These results indicate that these loci are simply highly expressed and particularly long hc-siRNA producing loci (Figure S3).

**Figure 3.**
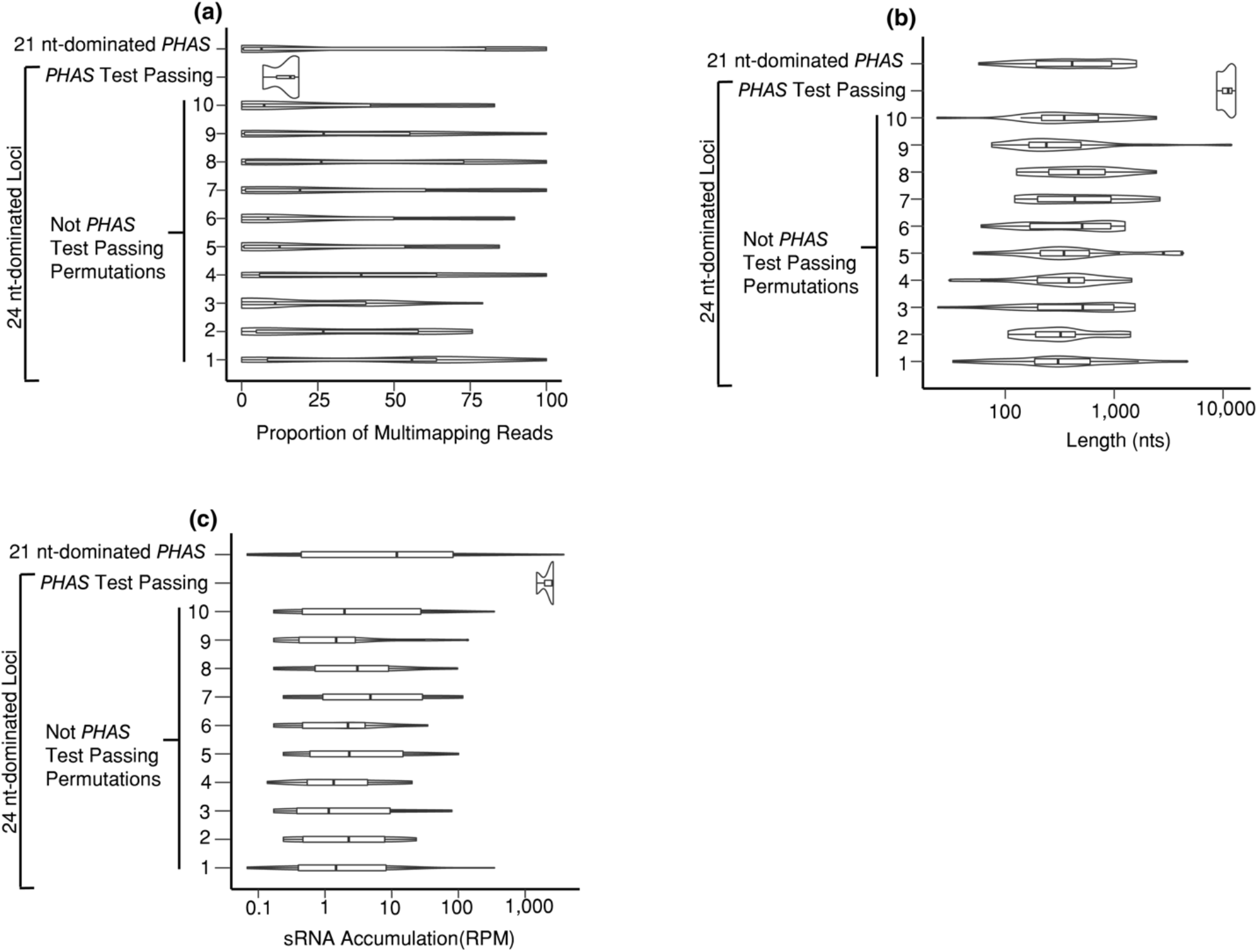
The three *PHAS*-Test passing small RNA loci have distinct properties compared to other 24 nt-dominated Loci. (a) The proportion of multi-mapped reads in three different types of small RNA loci. For 24 nt-dominated loci that were not called PHAS loci, ten cohorts comprising 20 randomly selected loci were used as controls. The width of the density plot shows the frequency. The inset boxes show medians (horizontal lines), the 1st-3rd quartile range (boxes), the 95% confidence of medians (notches), other data out to 1.5 times the interquartile range (whiskers) and outliers (dots). (b) Same as panel a except showing small RNA accumulation. (c) Same as panel a except showing small RNA locus length.

### A small number of *A. thaliana rdr6*-dependent, 24 nt-dominated siRNA loci exist, but are not phased

Canonical monocot 24 nt-dominated *PHAS* loci are produced in part through the biochemical activity of RDR6 (Zhai et al., 2015). We therefore used an alternative strategy to identify possible 24 nt *PHAS* loci by first identifying *rdr6*-dependent, 24 nt-dominated siRNA loci. We found 18 such loci (Figure S8; Dataset S1). None of these loci were consistently phased in the eight wild-type, inflorescence *A. thaliana* libraries (Table S2) tested according to any of the three algorithms we used (Dataset S3). We determined the genetic dependencies of these loci and found that these loci are down-regulated in *nrpd, nrpe*, and *rdr2* backgrounds (Figure 4). Furthermore, these loci either overlap transposable elements or are mostly found in otherwise intergenic regions (Figure S9a). As these are typical features of hc-siRNAs, it’s possible that hc-siRNAs simply erroneously placed at these loci. Again, we examined the proportion of ambiguously mapped reads in these *rdr6*-dependent, 24 nt-dominated loci compared to other 24 nt-dominated loci and found that these loci had low proportions of multi-mapping reads (Figure S9b). Our result argues against this hypothesis.

*rdr6*-dependent, 24 nt-dominated loci are strongly expressed relative to other 24 nt-dominated loci (Figure S9c). These *rdr6*-dependent loci have median expression level of nearly 100 RPM compared to the nearly 1 RPM median expression level of the other 24 nt-dominated loci (Figure S9c). This result is similar to the three *A. thaliana* loci that passed our *PHAS*-detection algorithms (Figure 3b). However, the lengths of the *rdr6*-dependent, 24 nt-dominated loci are similar to other 24 nt-dominated loci (Figure S9d). Overall while do find evidence for a small number of *rdr6*-dependent, 24 nt dominated siRNAs in *A. thaliana*, they are not phased, nor do they appear readily discernable from more typical hc-siRNA loci. Thus we find no evidence of 24 nt dominated *PHAS* loci in *A. thaliana*.

**Figure 4.**
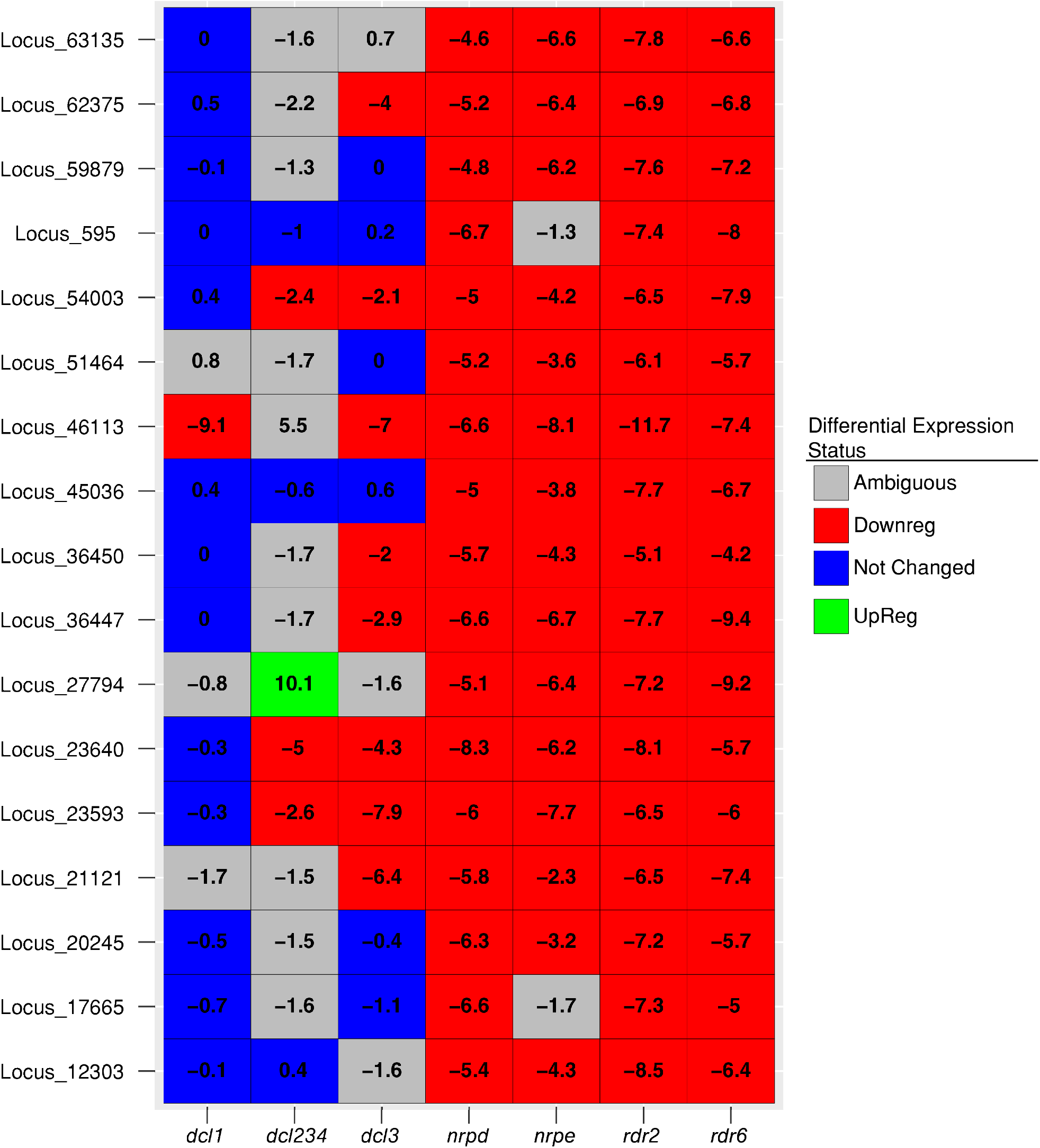
Accumulation of 24 nt-dominated, *rdr6*-dependent small RNA loci in different mutant backgrounds. Numbers represent the log2-transformed ratios of small RNA accumulation in the indicated genotypes over that in corresponding wild-type library as computed by DESeq2. The differential expression status was determined via DESeq2 at an FDR of 0.1.

### Erroneous detection of 24nt-domination *PHAS* loci occurs in other land plant species

We were interested in determining if erroneous annotation of *PHAS* loci was unique to *A. thaliana* small RNAs or if these results could be replicated in other species. We therefore interrogated publicly available *Brassica rapa, Cucumis sativus, Phaseolus vulgaris*, and *Solanum tuberosum* small RNA libraries (all eudicots) using the three algorithms, searching for putative 24 nt-dominated *PHAS* loci. We specifically chose these four species because they all lacked a potential *MIR2275* homolog (Figure 1). As miR2275 is the only miRNA known to trigger 24 nt-dominated phasiRNAs, any 24 nt-dominated loci called as *PHAS* loci in these species are likely false positives. Each species had 24 nt-dominated small RNAs that were misannotated as *PHAS* loci (Figure S10). We first determined where in the genome the 24 nt-dominated loci are encoded. Like in *A. thaliana*, 24 nt-dominated loci that passed the *PHAS*-detection algorithm seem to come predominantly from transposable elements. This was true in all the species examined except for *P. vulgaris* (Figure 5a). Curiously, the *PHAS*-test passing loci had a slightly higher proportion of ambiguously mapped reads compared to other 24 nt-dominated loci in *B. rapa*, *P. vulgaris*, and *S, tuberosum* (Figure 5b). For *C. sativus*, the proportion of multi-mapping reads in *PHAS*-test passing 24 nt-dominated loci was substantially higher than other 24 nt-dominated loci (Figure 5b). It’s still unlikely that multi-mapping reads contribute significantly to phasing at these loci as the proportion of ambiguously mapped to both types of 24 nt-dominated loci are similar in three of the four species tested (Figure 5b).

In the four species tested, the *PHAS*-test passing 24 nt-dominated loci had both significantly greater lengths (Figure 5c) and expression on average (Figure 5d). We observed this same trend in *A. thaliana* (Figure 3b-c), which suggests that long lengths and high expression levels of 24 nt-dominated loci are conducive to *PHAS* loci mis-annotations, even in other distantly related species.

**Figure 5.**
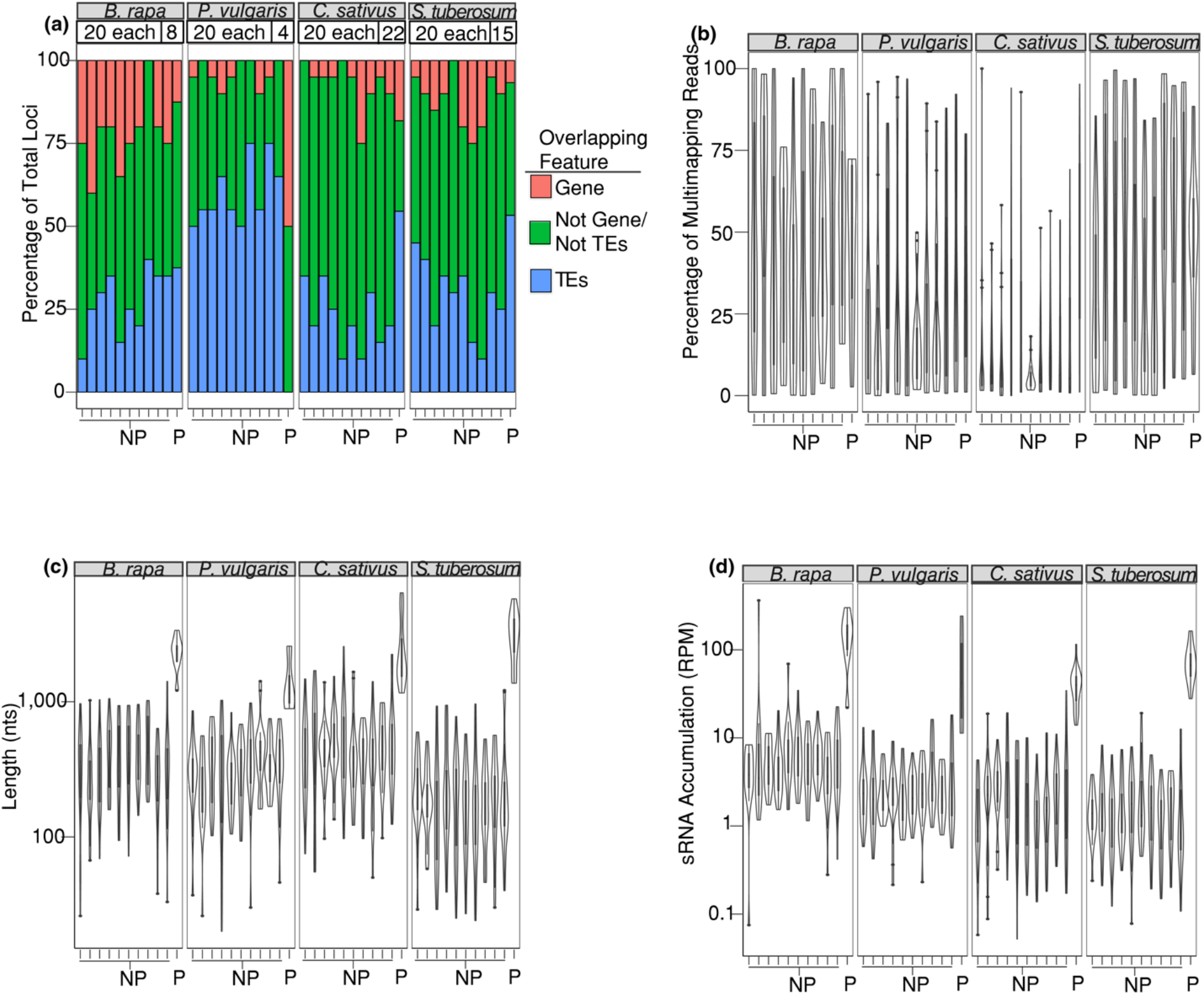
Four divergent species also contain 24 nt-dominated loci that passed the three *PHAS*-detecting algorithms. (a) Percentage of 24 nt-dominated Loci overlapping genes, transposable elements, or neither. The species name is shown in the top grey boxes. P: *PHAS*-test passing locus; NP: Not *PHAS*-test passing locus. For NP loci, ten cohorts of 20 randomly selected loci each were used as negative controls. The number of loci in each category is shown above the bar graphs. (b) Same as panel a except showing the proportion of multi-mapping reads. (c) Same as panel a except showing length of the small RNA locus. (d) Same as panel a except showing small RNA accumulation.

## Discussion

### *rdr6*-dependent, 24 nt-dominated loci in the *A. thaliana* genome

We found a handful of *rdr6*-dependent, 24 nt-dominated loci to be encoded in the *A. thaliana* genome. However, these loci have the same genetic dependencies as hc-siRNAs (Figure 4) and are frequently derived from the transposable elements (Figure S9a). The only other known *rdr6*-dependent, transposon-overlapping small RNA loci encoded in *A. thaliana* are epigenetically activated short interfering RNAs (easiRNAs) (Creasey et al., 2014). easiRNAs are derived from transcriptionally active transposable elements that are hypothesized to be targeted and cleaved by miRNAs (Creasey et al., 2014). Through the biochemical actions of RDR6 and DCL4, the miRNA cleavage product is converted into 21-22 nt double-stranded RNA molecules (Creasey et al., 2014). Furthermore, these easiRNAs are thought to direct initial repression of transposons (He et al., 2015). It’s possible that these *rdr6*-dependent, 24 nt-dominated loci could represent a transitory stage of hc-siRNA targeting, in which the genetic machinery of hc-siRNA are used but with a remaining initial dependency on RDR6.

### Possible losses of the *MIR2275*-generated phasing in eudicots

We examined all Phytozome (ver 12.1)-curated angiosperm genomes (Goodstein et al., 2012) for the presence of *MIR2275* homologs. *MIR2275* shows remarkable conservation in monocots (Figure 1), showing only loss in *Musa accuminata* and *Brachypodium distachyon*. This could be because these are aquatic plants with reduced morphology, and thus there is relaxed selective pressure for *MIR2275*. Another explanation is that the genome assemblies for these species could be incomplete. The lack of *MIR2275* in eudicots was more extensive (Figure 1), with no plants in the Brassicales clade containing a verified *MIR2275* homolog (Figure 1). Interestingly, *Aquilegia coerulea*, a basal eudicot (Sharma & Kramer, 2014), contained a *MIR2275* homolog (Figure 1; Figure S1) suggesting that the last common ancestor for eudicots may have contained *MIR2275*, and that the lack of detected putative *MIR2275* homologs in many eudicot plant species could be due to loss of *MIR2275*.

We only examined whether or not a possible *MIR2275* homolog could be detected by homology, if its predicted secondary structure is conducive to *MIRNA* processing (Figure S1), and in some species whether or not the putative *MIR2275* homolog was expressed in available small RNA libraries (Figure 1). We did not, however, determine if these species produce true 24 nt-dominated phasiRNAs. It’s possible that the potential *MIR2275* homolog in these species is an orphaned *MIRNA* or has taken on a new function. Furthermore, although some putative *MIR2275* homologs had *MIRNA*-like predicted hairpins, we note that they contain mismatches in the stem of the secondary structure which may hinder miRNA biogenesis (Figure S1). Further research is necessary to determine if these species truly encode 24 nt-dominated *PHAS* loci.

### The difficulties of annotating 24 nt-dominated *PHAS* loci

The discovery of 24 nt-dominated *PHAS* loci in maize (Zhai et al., 2015), rice (Song et al., 2008), asparagus, daylily, and lily (Kakrana et al., 2018) opened up the possibility that these loci exist in other species, even distantly-related eudicots. However, 24 nt-dominated hc-siRNA loci are widespread in land plant genomes (Ghildiyal & Zamore, 2009). This is true even in *A. thaliana*, although 24 nt sRNAs are thought to repress transposable elements (Matzke et al., 2009) and only about 20-30% of the *A. thaliana* genome consists of transposon/repetitive elements (Barakat et al., 1998). In contrast, other plant genomes consist of as much as 80% transposons/repetitive elements (Springer et al., 2009). The sheer number of 24 nt-dominated sRNA loci could mean that several of them could meet various annotation criteria simply by chance; this is something we have previously noted to occur in natural antisense siRNA annotation (Polydore & Axtell, 2018).

The supposed 24 nt-dominated *PHAS* loci examined in this study consistently showed higher levels of expression than other 24 nt-dominated sRNA loci and derived from very long loci. This isn’t particularly surprising because the accumulation of reads (in- and out-of-phase) is a factor in most *PHAS*-detection algorithms (Figure S3). However, it seems that particularly highly expressed 24 nt-dominated small RNAs are able to consistently pass *PHAS*-detection algorithms because of this. It’s possible that the sheer number of reads produced at these loci means that these loci produce enough “in-phase” reads by chance to score highly in *PHAS*-detecting algorithms.

### Annotating 24 nt-dominated *PHAS* loci in the future

Our study demonstrates that when annotating novel 24 nt-dominated *PHAS*, more than utilizing *PHAS*-detecting algorithms is necessary for robust annotation. First, examining different available mutants, especially of genes involved in small RNA biogenesis, can be critical in determining type of small RNA in question (Figure 2c). All types of phasiRNAs are known to be reliant on the biochemical activity of RDR6 and Pol II (Cuperus et al., 2010; Song et al., 2008; Zhai et al., 2015). While it’s entirely possible that non-canonical phasiRNAs that are reliant on RDR2 and Pol IV may be described eventually, such a study must verify with robust methodologies that the *PHAS* loci aren’t false positives due to the sheer number of Pol IV/RDR2 –dependent reads produced in the land plant genome.

We searched for *MIR2275* homologs because miR2275 is the only miRNA known to trigger 24 nt phasiRNAs. If a species lacks a *MIR2275* homolog, then one should be skeptical of any 24 nt-dominated small RNA locus that is annotated as *PHAS* locus in that specie. An organism producing a mature miR2275 small RNA homolog is not sufficient to show that 24 nt-dominated *PHAS* loci are produced in that species. One should also take care to ascertain if any 24 nt-dominated small RNA locus called as *PHAS* loci also contains a miR2275 target site that is “in phase” with the phasiRNAs produced from the transcript. miR2275 is also very specifically expressed in the tapetum of male floral tissue. In asparagus, 24 nt-dominated phasiRNAs have also been shown to be expressed in female floral tissue. However, 24 nt-dominated phasiRNAs have not been demonstrated to be expressed outside of reproductive tissue as of yet. Therefore, any 24 nt-dominated *PHAS* loci annotated in libraries not produced from reproductive tissues are more likely to be mis-annotations.

Despite the fact that several small RNA loci in *A. thaliana* are able to consistently pass the *PHAS*-detecting algorithms in different libraries, reproducibility among distinct small RNA libraries is of utmost importance. It could also be useful to employ several different *PHAS*-test algorithms as well. In *A. thaliana*, three loci were able to consistently pass our *PHAS*-detecting algorithms (Figure S5), but the number of loci that each algorithm detected on its own was generally much higher (Figure 2a). Had we not used stringent *PHAS*-detecting methods, it would’ve been plausible to assume we had found 24 nt-dominated *PHAS* loci in *A. thaliana* based on the number of loci that consistently pass alone. Utilizing multiple small RNA libraries should become easier to accomplish as more and more libraries in different treatments/genotypes become available for socio-economically relevant species. Certain *PHAS*-detecting programs, such as *PHASIS* (Kakrana et al., 2017), automatically evaluate potential small RNA loci in different libraries individually before merging the results of each library.

It’s not always possible to determine the small RNA that targets the phasiRNA precursor transcript. Indeed, several reads may be predicted to target a certain transcript by chance due to the sheer number of unique reads in a small RNA library. However, due to hydrolysis of the precursor transcript following small RNA-mediated targeting, phasiRNAs have well-defined termini from which they are produced (Axtell et al., 2006; Cuperus et al., 2010). Therefore, a true *PHAS* loci should have a large proportion of its reads reproducibly falling into a particular phase register. This is what we observed with well-characterized *PHAS* locus*, TAS2*, but not with the three loci that pass our rigorous *PHAS*-annotation regime in *A. thaliana* (Figure S6).

As 21-22 nt-dominated *PHAS* loci are far more common in land plants, especially outside of the monocots, one can conservatively limit their discovery of new *PHAS* loci to 21-22 nt-dominated small RNAs and not employ as rigorous methodology as the ones used in this study. However, one should still employ post *PHAS*-discovery quality controls to ensure these 21-22 nt-dominated *PHAS* loci are genuine (such as determining if the predominant phase registers are reproducibly dominant at these loci (Figure S6)). 24 nt-dominated *PHAS* loci on the other hand seem to be have undergone loss in many angiosperms. However, even in the species in which they are conserved, they seem to play very specific, reproductive-associated roles as evidenced by their expression patterns. Great caution should be used for annotating 24 nt-dominated *PHAS* loci in the future.

## Materials and Methods

### Finding potential *MIR2275* homologs in angiosperms

The Phytozome (ver 12.1)-curated angiosperm genomes sequences were downloaded. The mature *O. sativa* miR2275a sequence was downloaded from miRBase (ver 21.) (Griffiths-Jones et al., 2008) and searched against all the other genomes using Bowtie v1.0 (Langmead et al., 2009) allowing for two mismatches.

In order to determine the predicted secondary structures of the Bowtie results of interest, the sequences corresponding to the Bowtie result, plus 200 nucleotides upstream and downstream, was extracted from the specie’s genome. The secondary structures of the sequences were predicted using the mFOLD web server (Zuker, 2003) and visually inspected to determine if the sequence formed a hairpin structure consistent with accepted norms for miRNA biogenesis (Axtell & Meyers, 2018). The sequences were aligned using ClustalX ver. 2 with default parameters (Larkin et al., 2007).

For those species for which publicly available small RNA-seq data existed, we downloaded, trimmed, and merged the small RNA libraries. The merged libraries were collapsed to non-redundant reads and investigated using CD-HIT (ver 4.6.8) using the options -n 4, -d 0, and -g 1. The *O. sativa* miR2275a sequence was used as a query.

### Determination of potential 24 nt-dominated *PHAS* loci in different wild-type, inflorescence libraries

Wild-type biological triplicate small RNA libraries (GSE105262) (Polydore & Axtell, 2018) were merged and aligned against the *A. thaliana* (TAIR 10) genome using ShortStack (ver 3.8) (Johnson et al., 2016) using 27 known *A. thaliana PHAS* loci as a query file (Table S1). Three distinct *PHAS* –detecting algorithms (Dotto et al., 2014; Johnson et al., 2016; Zheng et al., 2014) were used to determine phase scores in the merged run. These scores were used as the basis to determine cutoffs for calling significantly phased loci (Figures S3-S4). Phase scores of each known *PHAS* locus for each of the three algorithms in the eight wild-type libraries used is listed in Dataset S4. Note that we didn’t use the multiple testing correction for *PHAS* loci p-values as done in Dotto et al. 2014 as we wished to test the three algorithms against each other, and the algorithms that yield phase scores couldn’t be adjusted for multiple testing.

Wild-type and *rdr1-1/2-1/6-15* (*rdr1/2/6*) triple mutant small RNA libraries (GSE105262) (Polydore & Axtell, 2018) were aligned against the *A. thaliana* (TAIR 10) genome using ShortStack (ver 3.2) with option –pad 75 and option -min_cov 0.5rpm (Table S2). With these settings, small RNA loci are found as follows: All distinct genomic intervals containing one or more primary sRNA-seq alignments within 75 nts of each other were obtained, and then filtered to remove loci where the total sRNA-seq abundance with a locus was less than 0.5 reads per million. This produced a final set of distinct, non-overlapping small RNA loci. Differential expression to determine down-regulated loci was performed as previously described (Polydore & Axtell, 2018). Down-regulated, 24 nt-dominated small RNA loci were catalogued into a list. Eight wild-type, inflorescence small RNA libraries (Table S2) were run individually against the *A. thaliana* (TAIR 10) genome utilizing ShortStack (ver 3.8.1) using the results of our wild-type and *rdr1/2/6* libraries run as a query file. The phase scores of loci corresponding to the down-regulated, 24 nt-dominated small RNAs loci were evaluated using the binary sequence alignment (BAM)-formatted alignments from each run. ShortStack and an in-house Python script was used to perform the three phase score calculations. For *B. rapa*, *C. sativus*, *P. vulgaris*, and *S. tuberosum*, the previous Shortstack merged small RNA alignments were analyzed in the same way.

For the four other species besides *A. thaliana*, we downloaded publicly available small RNA libraries (Dataset S2) for *Brassica rapa*, *Cucumis sativus*, *Phaseolus vulgaris*, and *Solanum tuberosum* and merged and aligned them against their respective genomes using ShortStack (ver 3.8.1) with default options. The genomes used were ver 1.0 for *B. rapa* (Wang et al., 2011), ver 1.0 for *P. vulgaris* (Schmutz et al., 2014), ver. 2 for *C. sativus* (Huang et al., 2009), and dm_v404 for *S. tuberosum* (Hardigan et al., 2016).

### AGO Immunoprecipitation, genetic dependency, and properties of loci analyses

Calculating the lengths, proportion of multi-mapping reads, small RNA expression levels (in Reads Per Million (RPM)), determining the genetic dependencies, and the AGO enrichments of various loci were performed as previously described (Polydore & Axtell, 2018). In order to compare the properties of 24 nt-dominated loci that passed the *PHAS*-detection algorithms to those that didn’t, 10 subsets of 20 loci were randomly selected from the 24 nt-dominated loci that didn’t pass the *PHAS*-detection algorithms with replacement.

### Examining *PHAS*-test passing loci for potential miRNA targeting

Sequences corresponding to the *A. thaliana* loci that consistently passed the *PHAS*-detection algorithms plus 200 base pairs up-stream and downs-stream were extracted from the *A. thaliana* genome. Mature miRNA sequences were downloaded from miRBase (ver 21) and aligned against these sequences using the Generic Small RNA-Transcriptome Aligner. A miRNA was considered to be potentially targeting a sequence if it aligned with an Allen et. al. score of 3 or less.

## Acknowledgements

This research was funded by NSF Award 1339207 to MJA, the NIH Bioinformatics and Statistics (CBIOS) Predoctoral Training Program (1T32GM102057-0A1) to SP, and the Henry W. Popp Graduate Assistantship Fund to SP.

## Supplementary Figures

**Figure S1.**
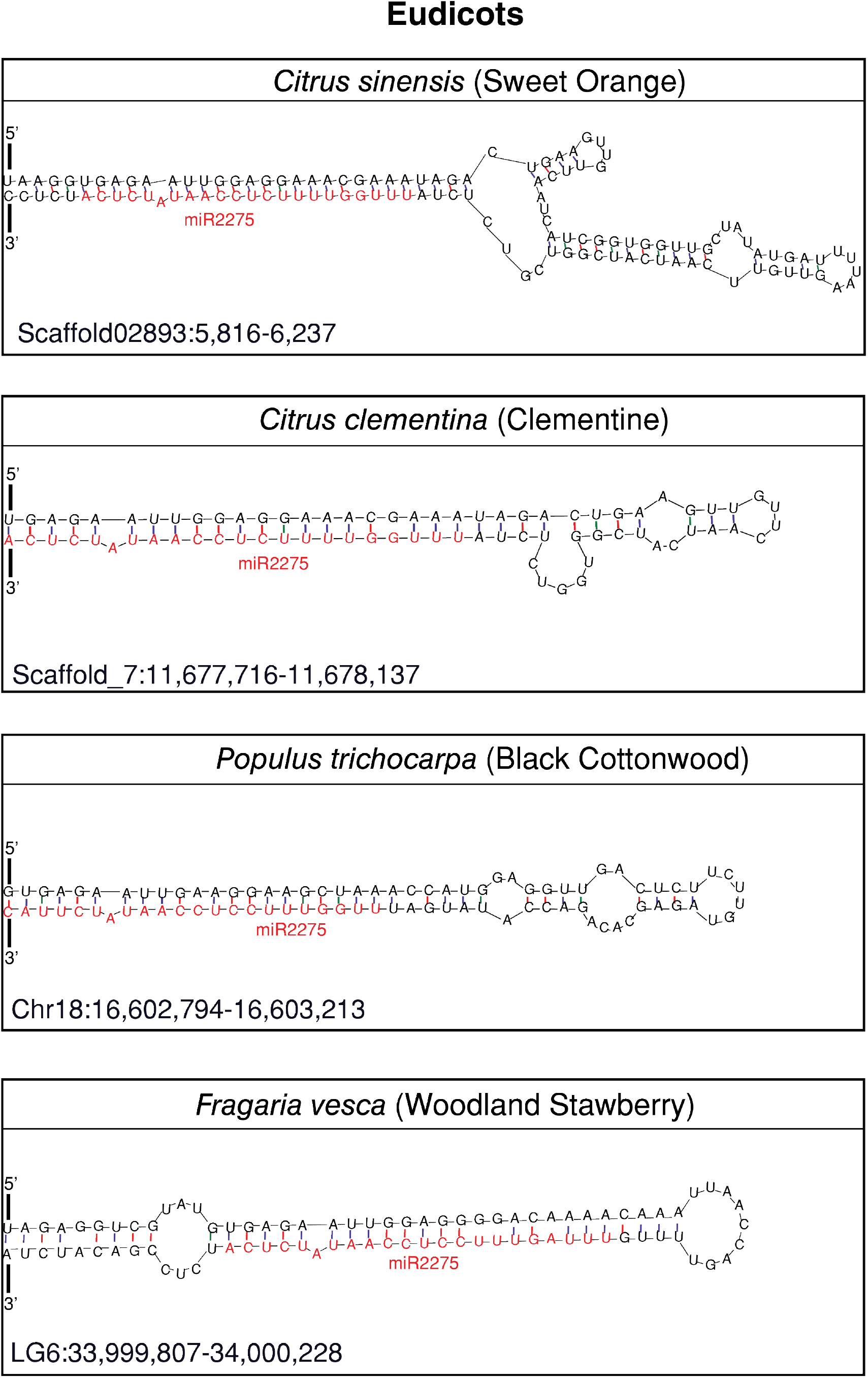
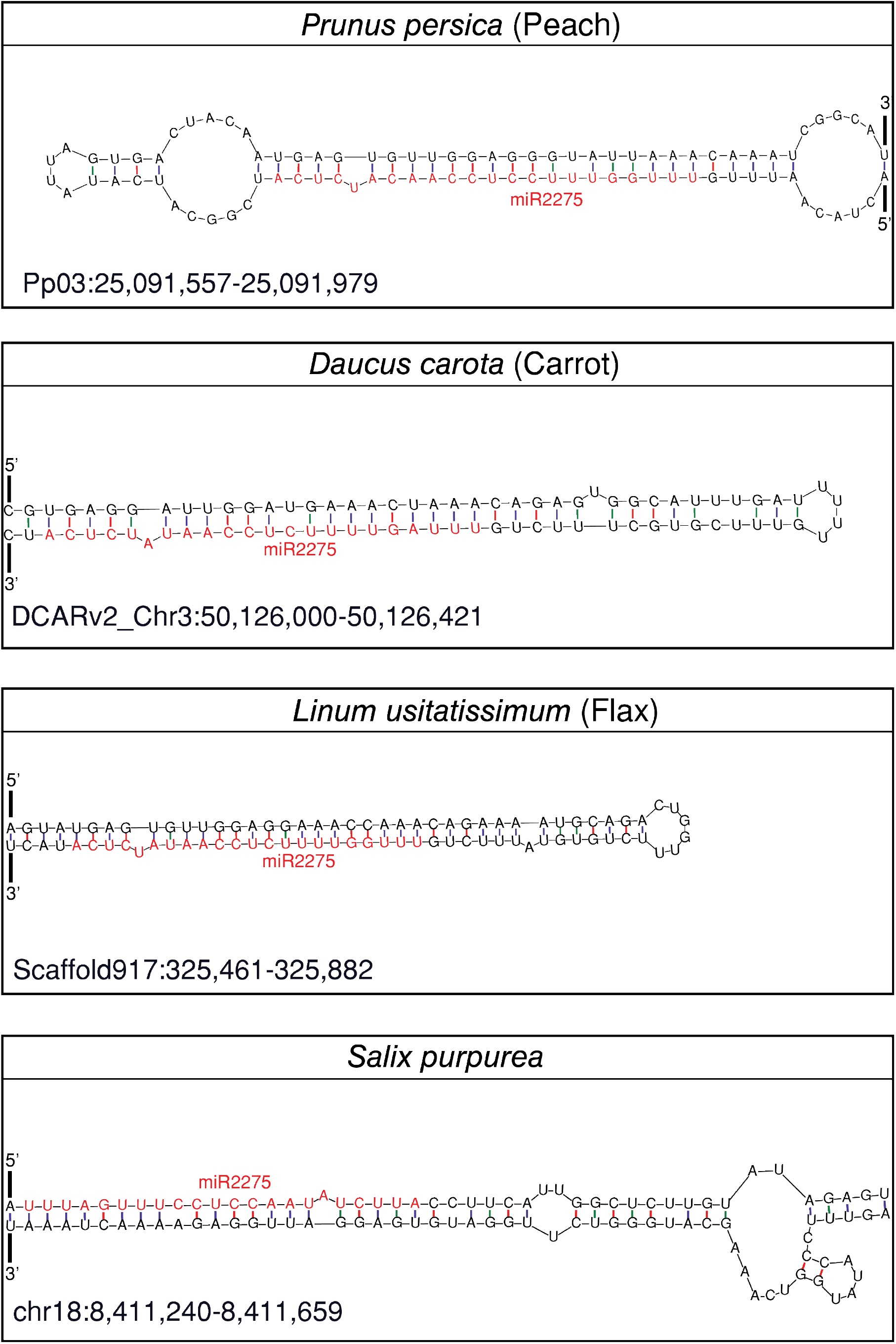
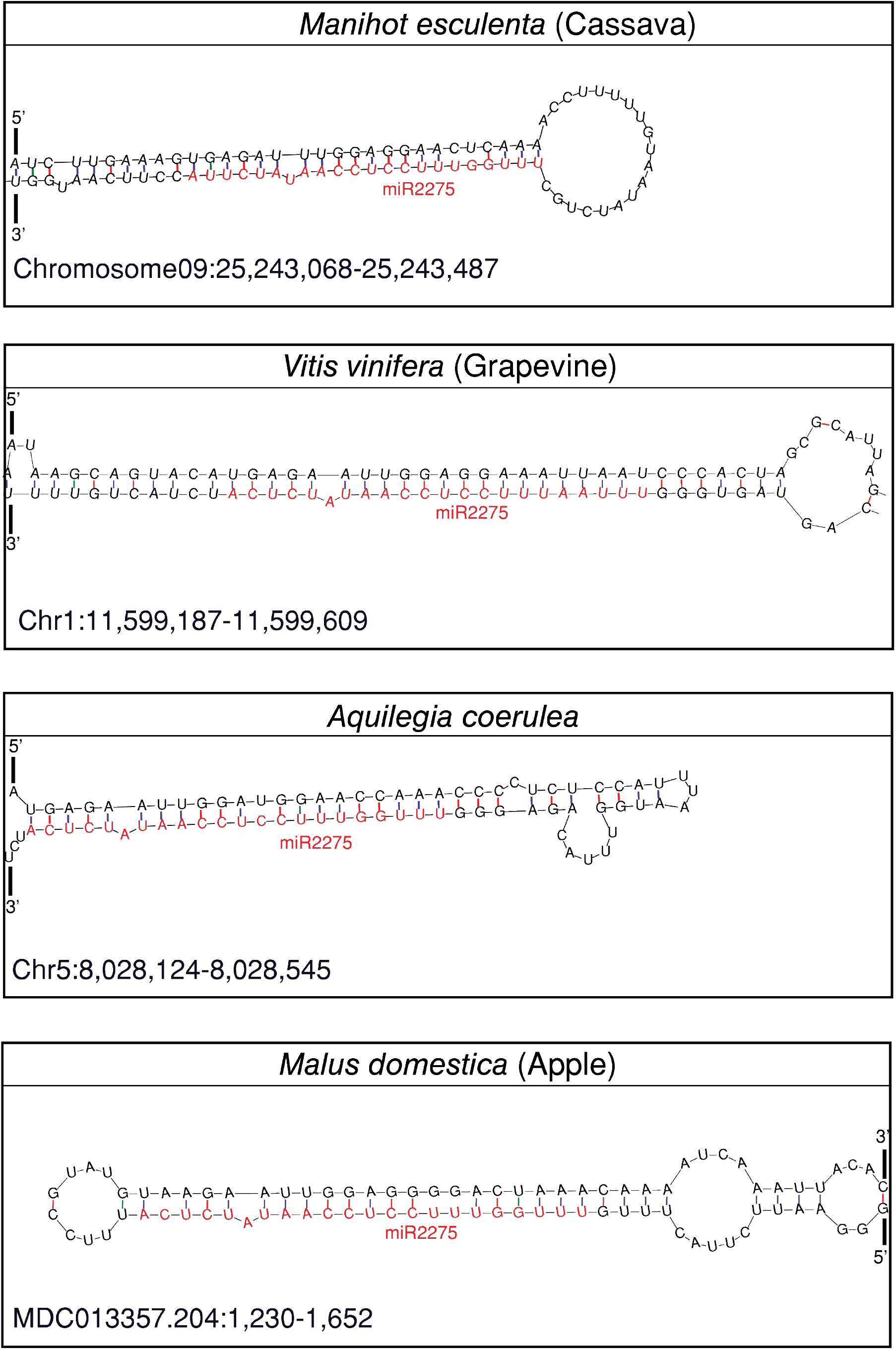
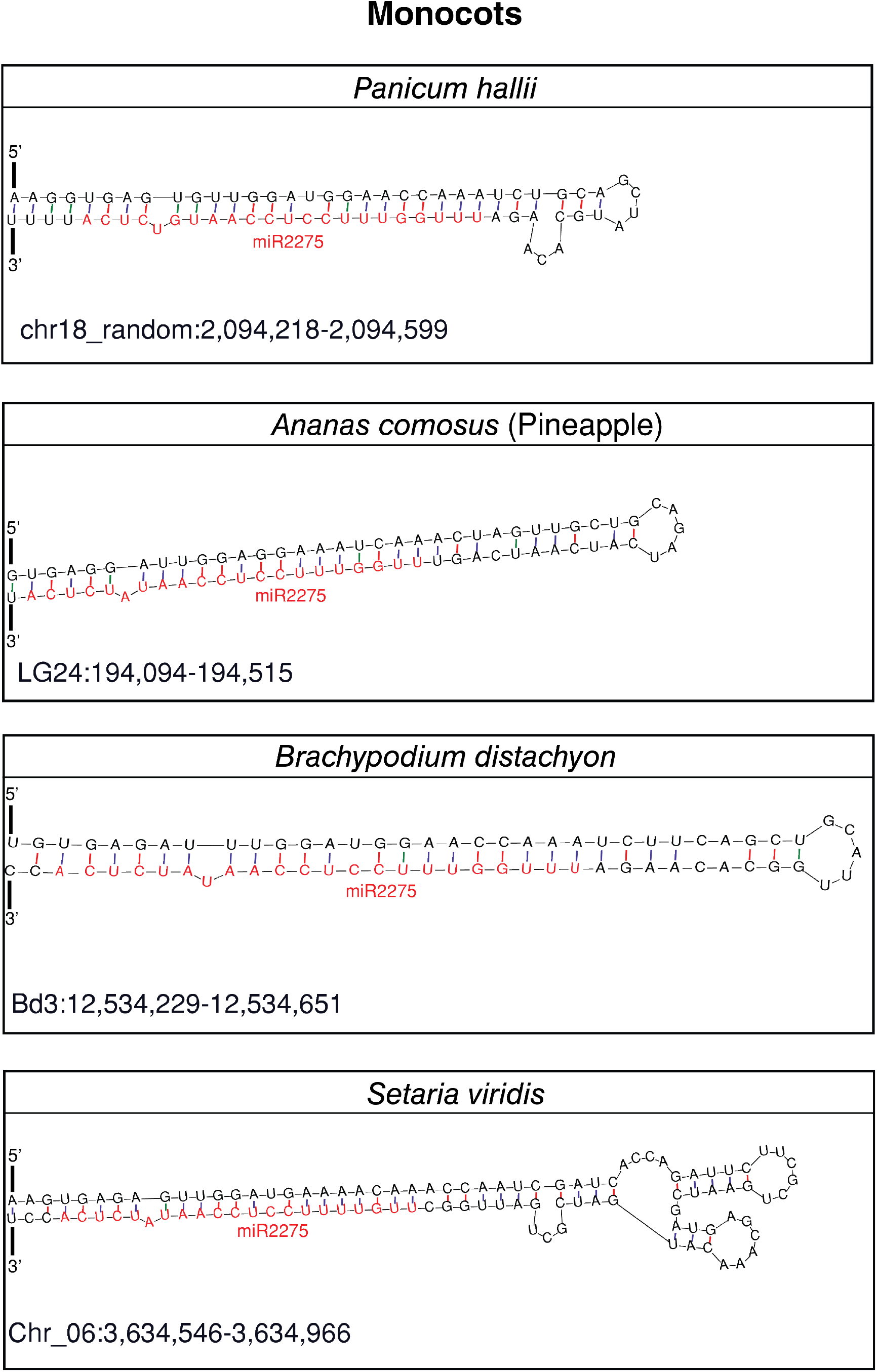
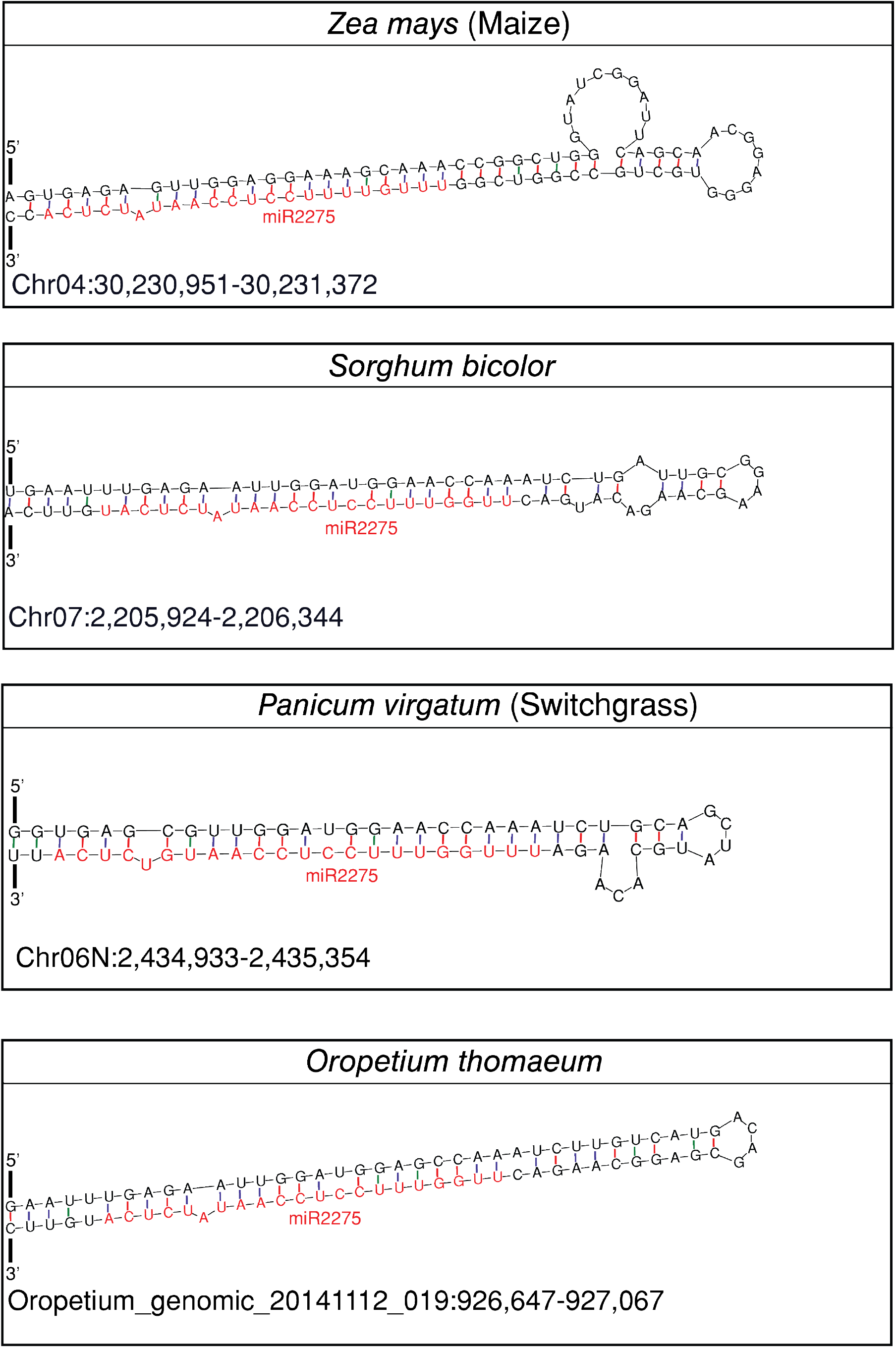
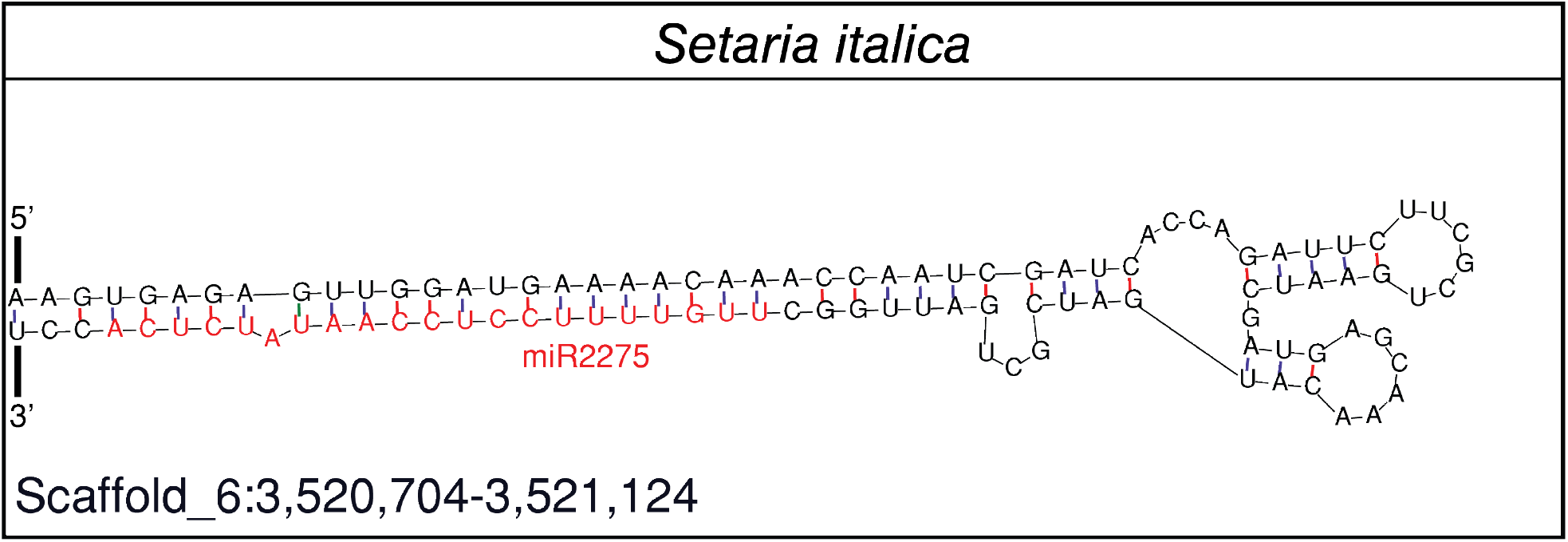
The RNA secondary structure of Bowtie-validated *MIR2275* homologs. Secondary structures of loci were predicted via mFOLD. If the potential *MIR2275* homolog formed a stem-and-loop structure, it is shown above. The red nucleotides denote the mature *MIR2275* homolog in the secondary structure.

**Figure S2.**
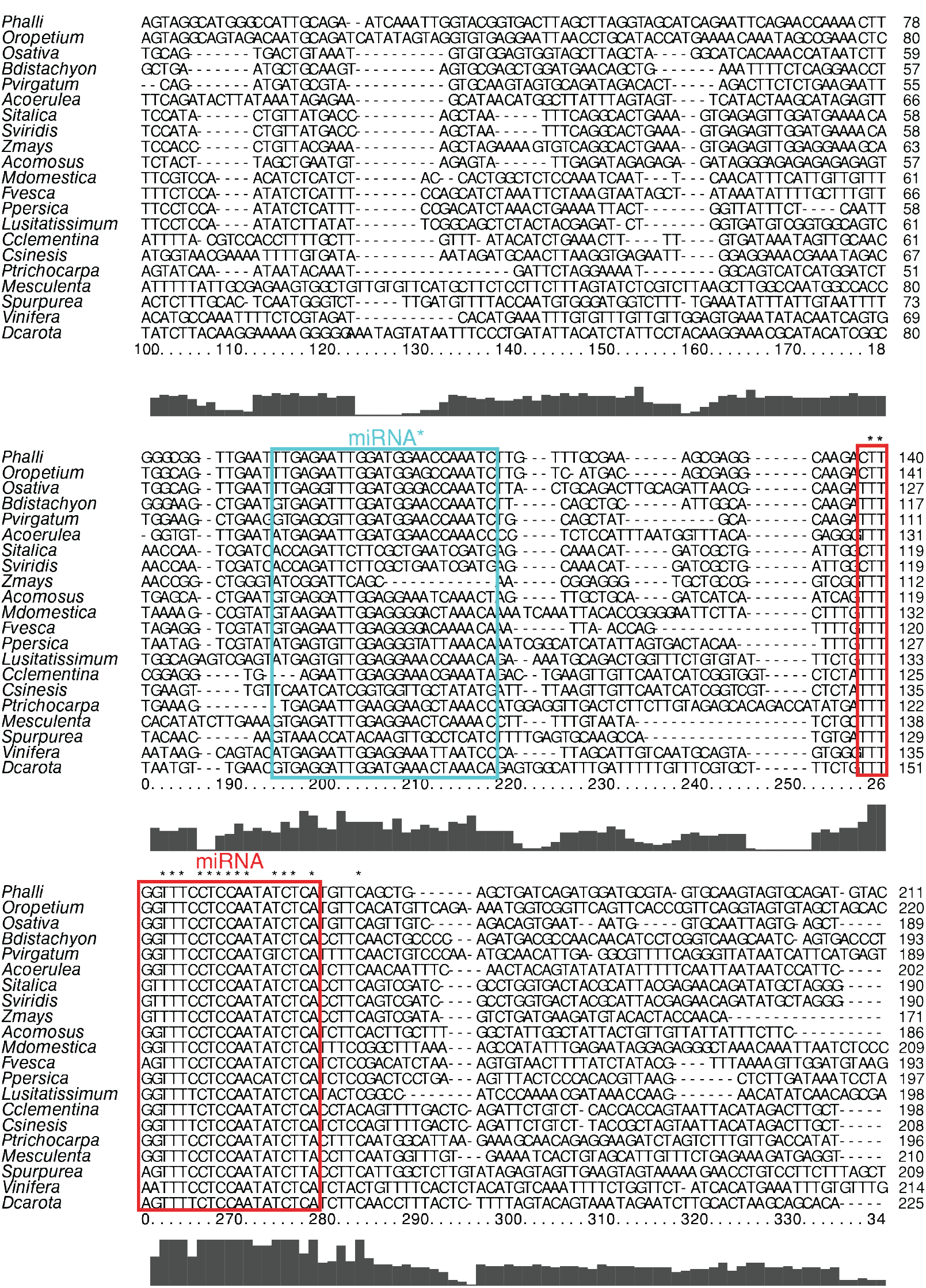
Alignment of *MIR2275* homologs. Sequences corresponding to miR2275 wih flanking 200 bp islands were extracted from genomes of the above-listed genomes and aligned via ClustalX2. Only the 100^th^ to 340^th^ nucleotide of each sequence (including gaps) are shown. miR2275 and miR2275* in each species are shown.

**Figure S3.**
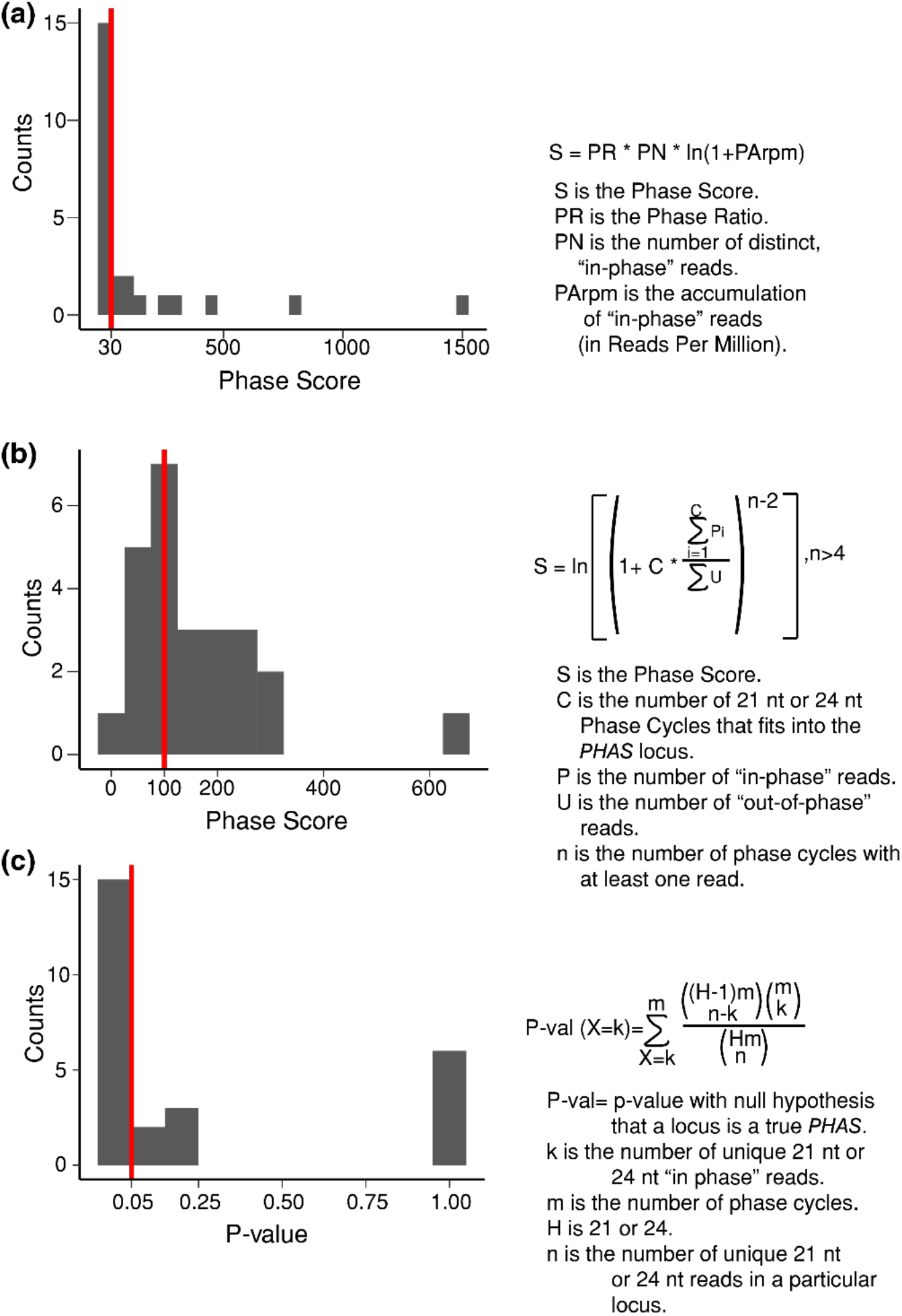
Determination of phase score cutoffs using three different algorithms. (a) Small RNA accumulation at 27 known 21 nt *PHAS* loci (Table S1) was analyzed using the *PHAS*-Test algorithm from Guo et al. (2015), which is used by ShortStack. Variables and formula are shown. Red line represents the cut-off used in this study to determine if a locus was phased or not; scores above the red line were considered ’phased’. (b) Same as in panel a except for the *PHAS*-Test algorithm from Dotto et al. (2014). (c) Same as in panel a except for the *PHAS*-Test algorithm from Zheng et al. (2014); scores below the red-line were considered ‘phased’.

**Figure S4.**
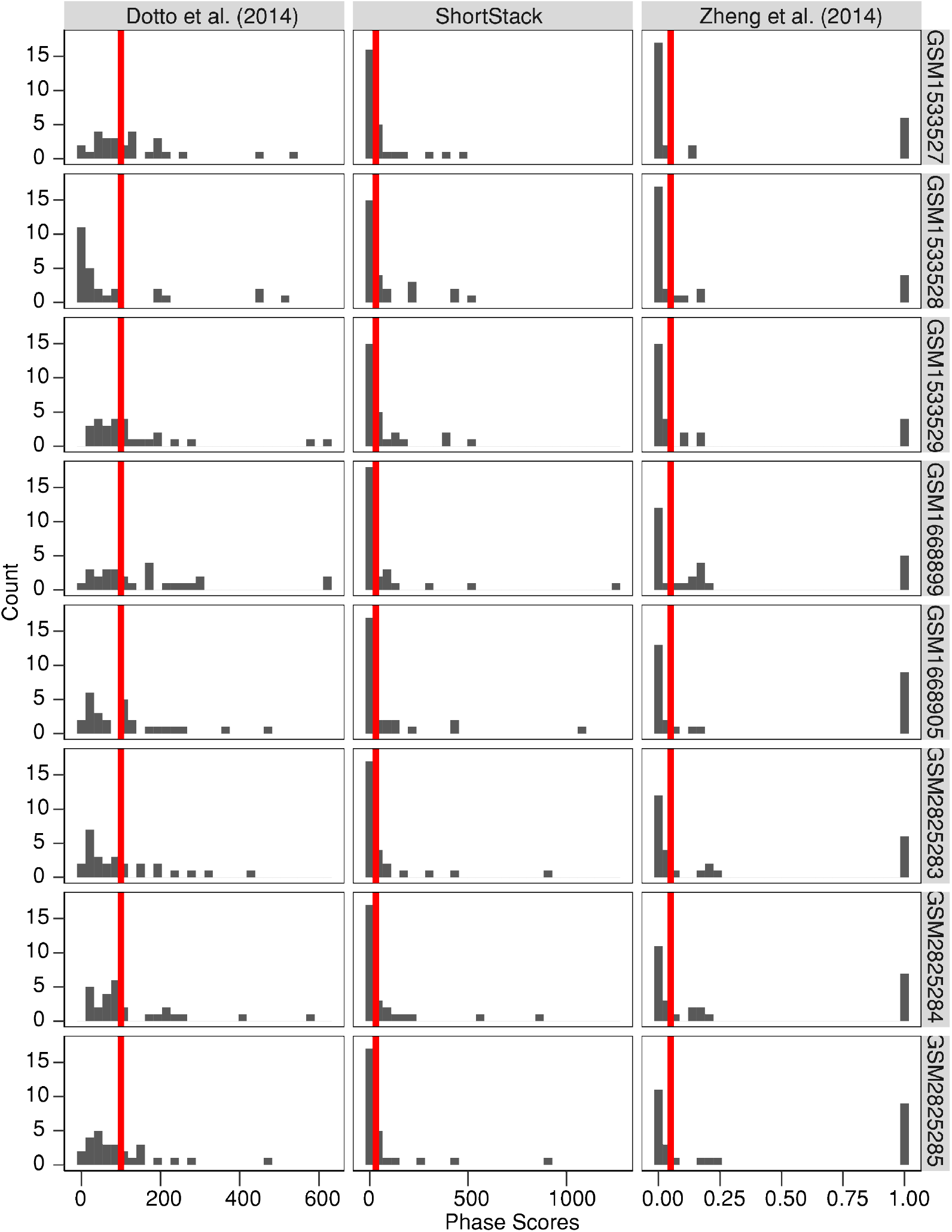
Phase scores of 27 previously known *A. thaliana* phased siRNA loci from the indicated algorithms. Cutoff values for called ’phasing’ in our study are shown in red.

**Figure S5.**
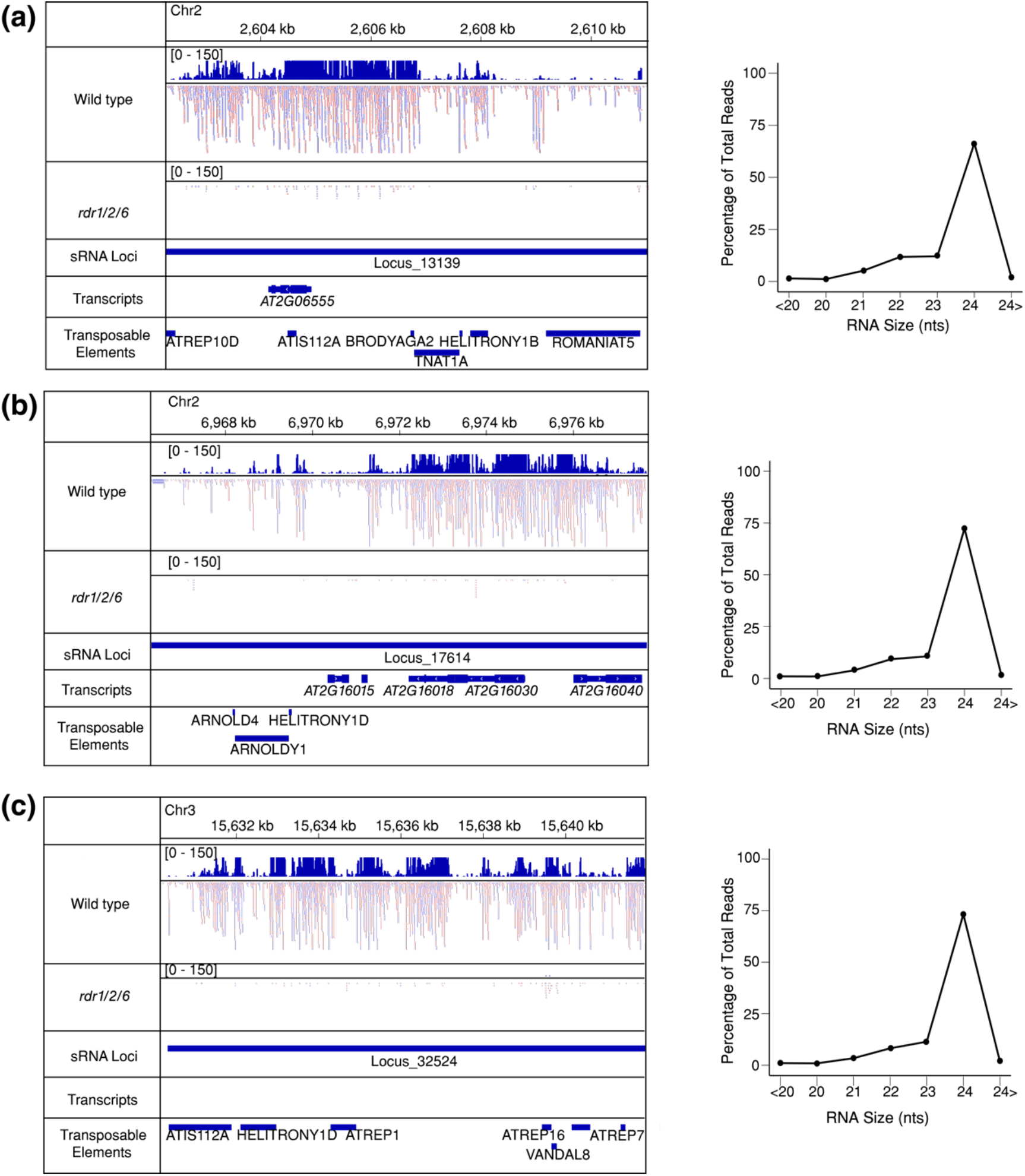
Genome browser snapshots of the three *Arabidopsis thaliana PHAS*-Test passing loci. (a) Alignments and read size distribution of Locus_13139. Red reads represent positive-strand mapped genes, while blue reads are those that mapped to the negative strand. Numbers in the brackets are the range of coverage shown in Reads per Million. (b) Same as a, except for Locus_17614. (c) Same as a, except for Locus_32524.

**Figure S6.**
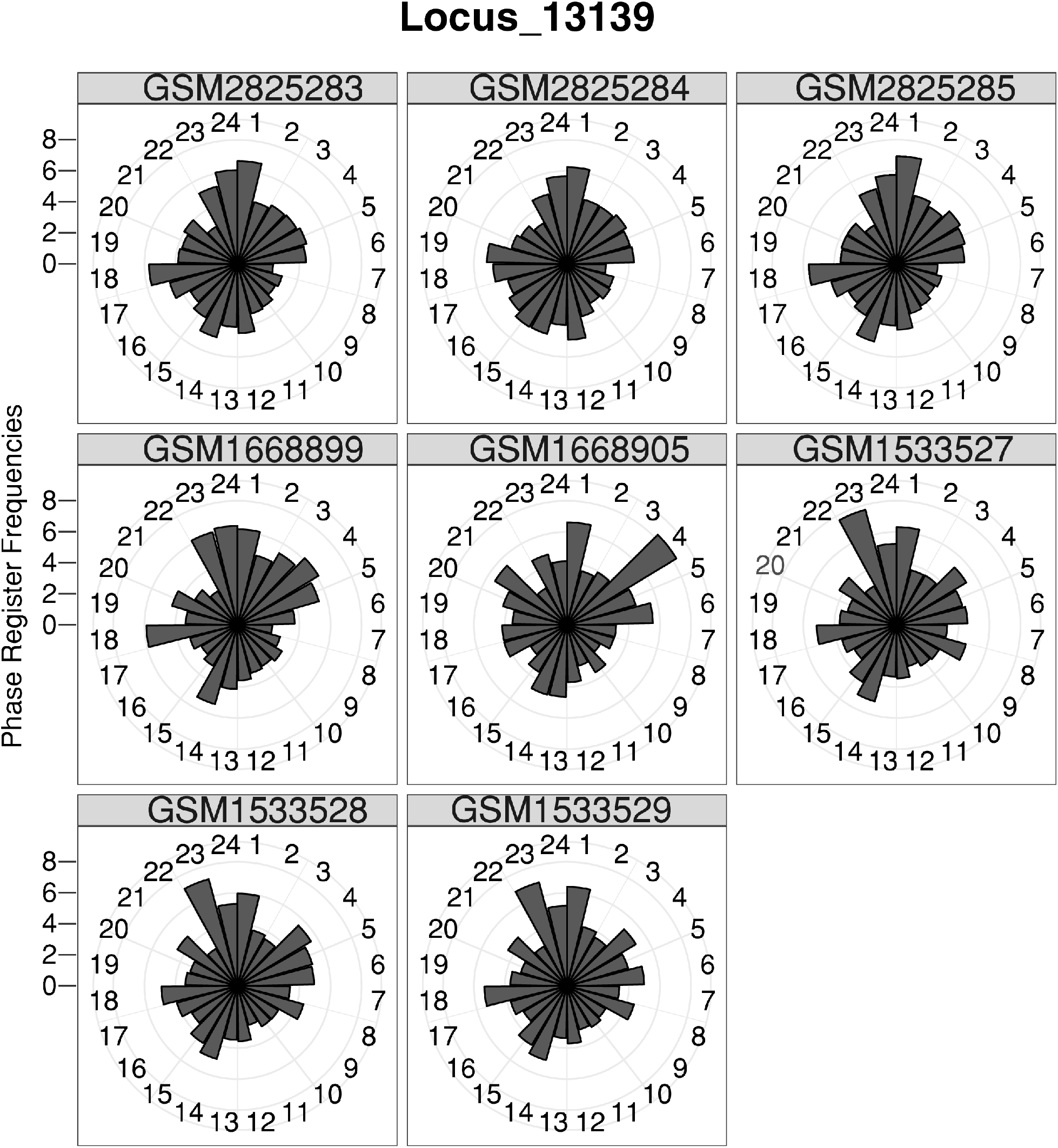
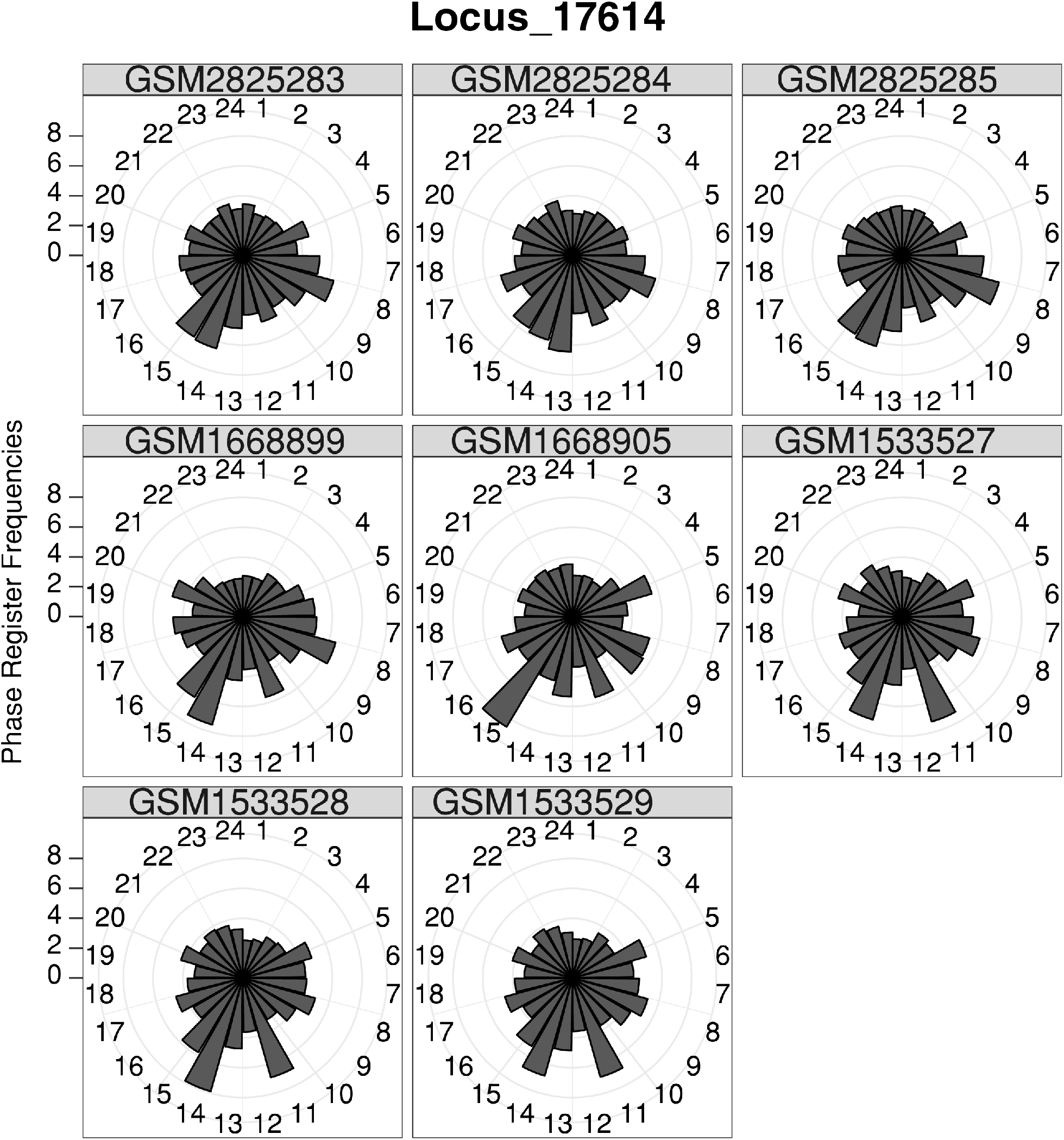
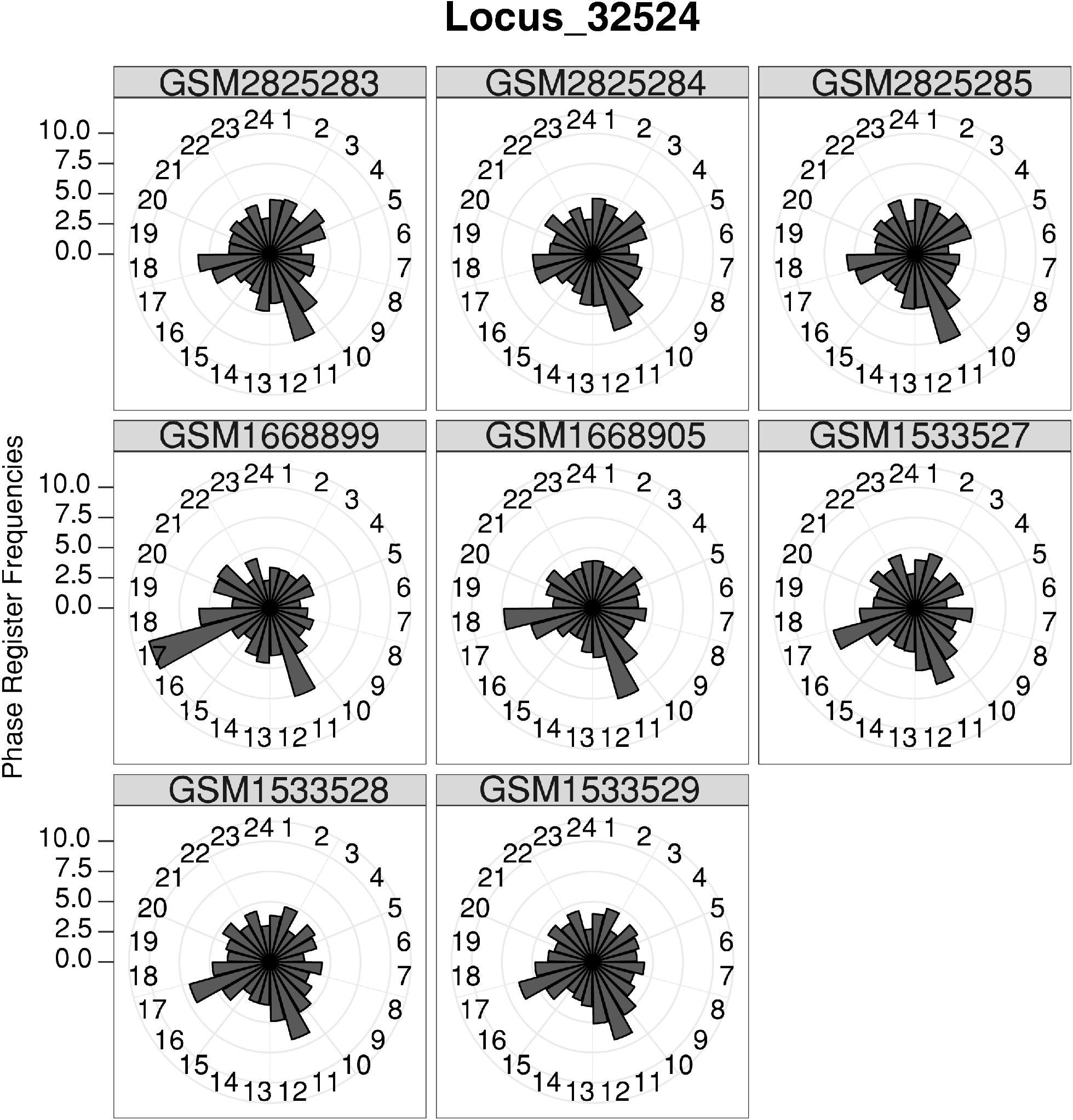
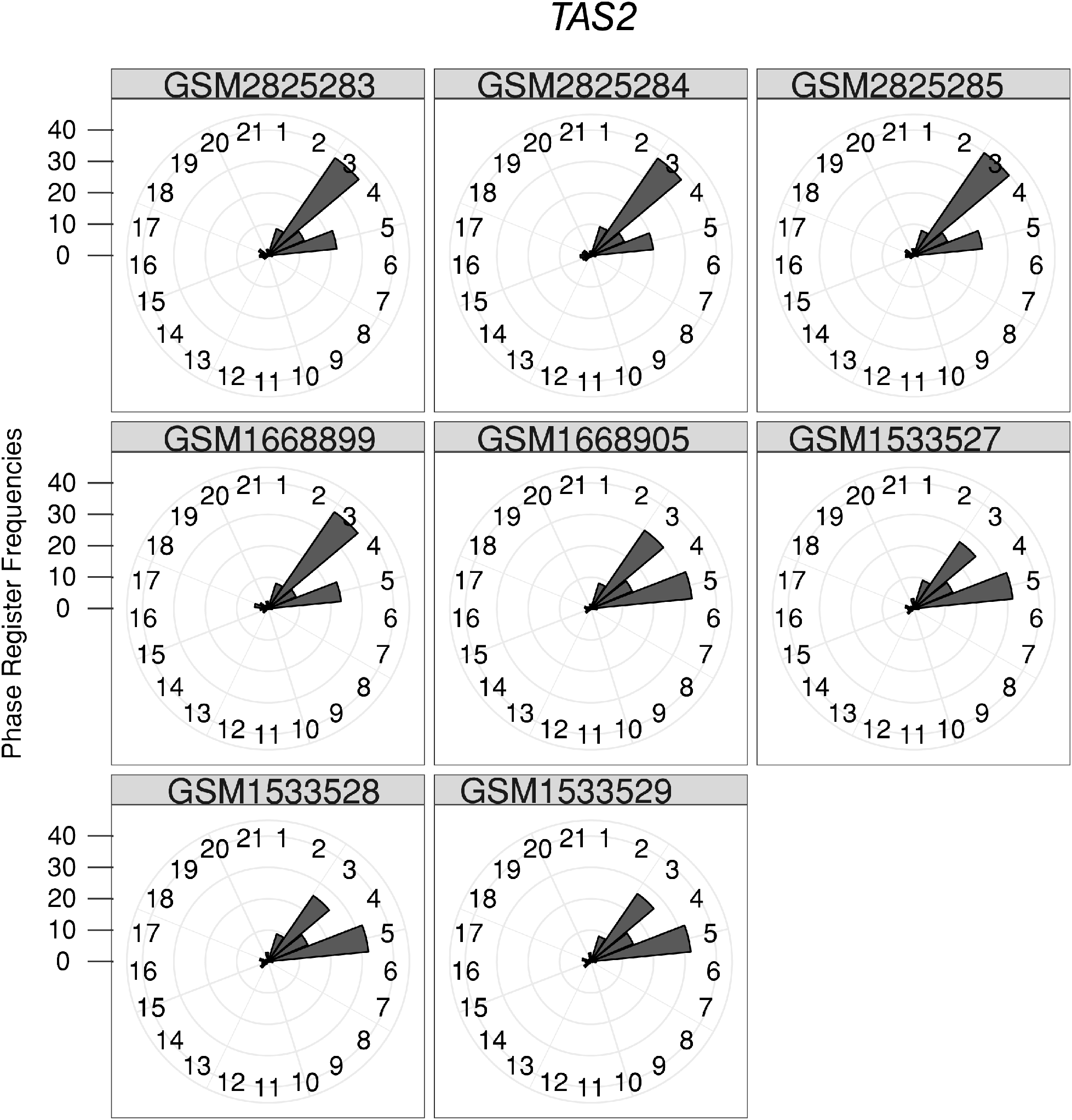
The three *PHAS*-Test passing loci have inconsistent phase register frequencies. Frequencies of phase registers where calculated in eight publicly available wild-type inflorescence libraries from *A. thaliana* (the accession numbers are listed in the grey boxes). *TAS2* is shown for comparison. Phase register frequencies are calculated via the following formula: (number of reads “in phase”/ total number of reads at locus)*100.

**Figure S7.**
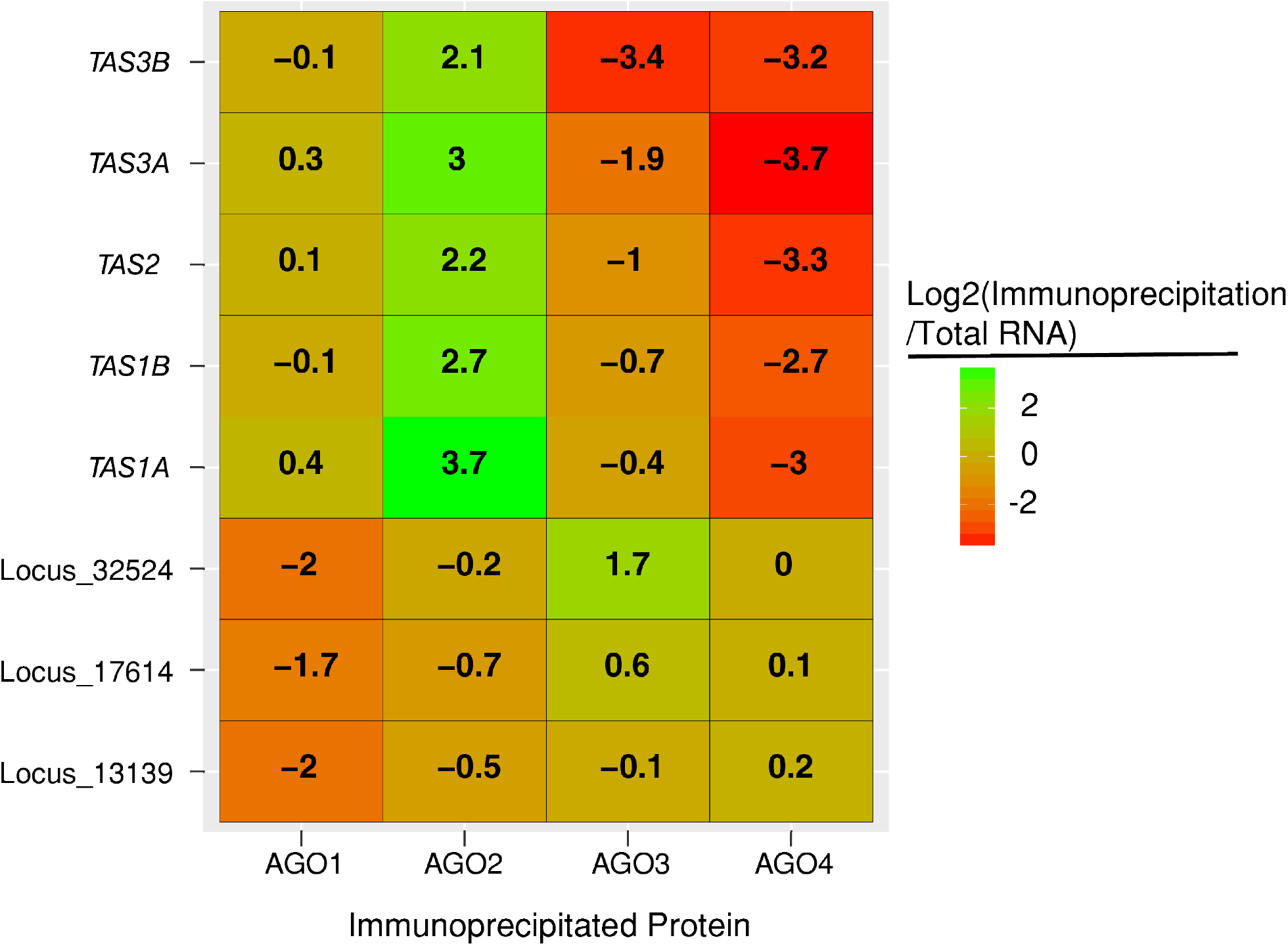
The three putative 24 nt *PHAS* loci show AGO-loading profiles that are distinct from *TAS* loci. Small RNAs from immunoprecipitation protein were aligned to the *A. thaliana* (TAIR10) genome. Numbers indicate the ratio of sRNA accumulation between immunoprecipitated and total libraries (in RPMs).

**Figure S8.**
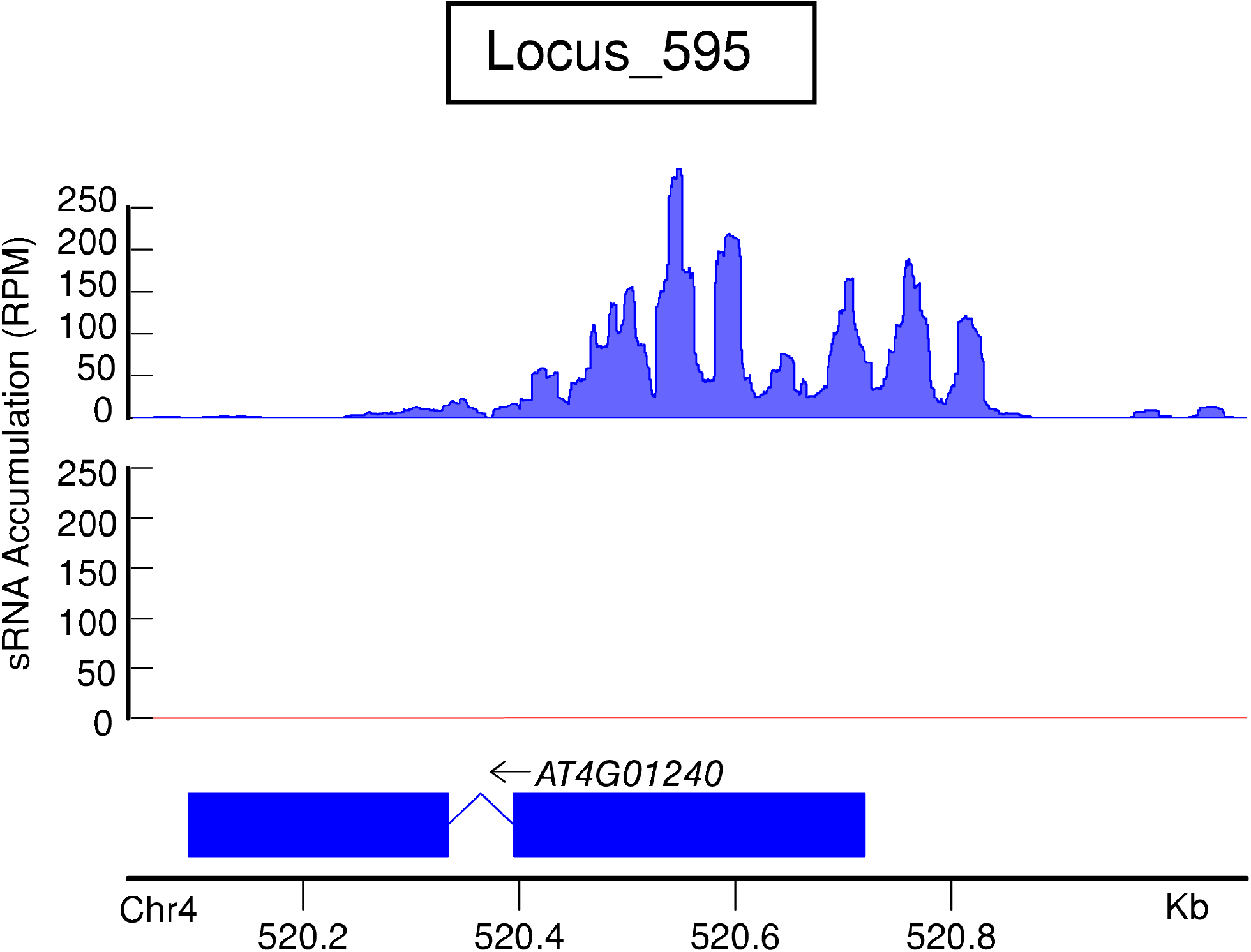
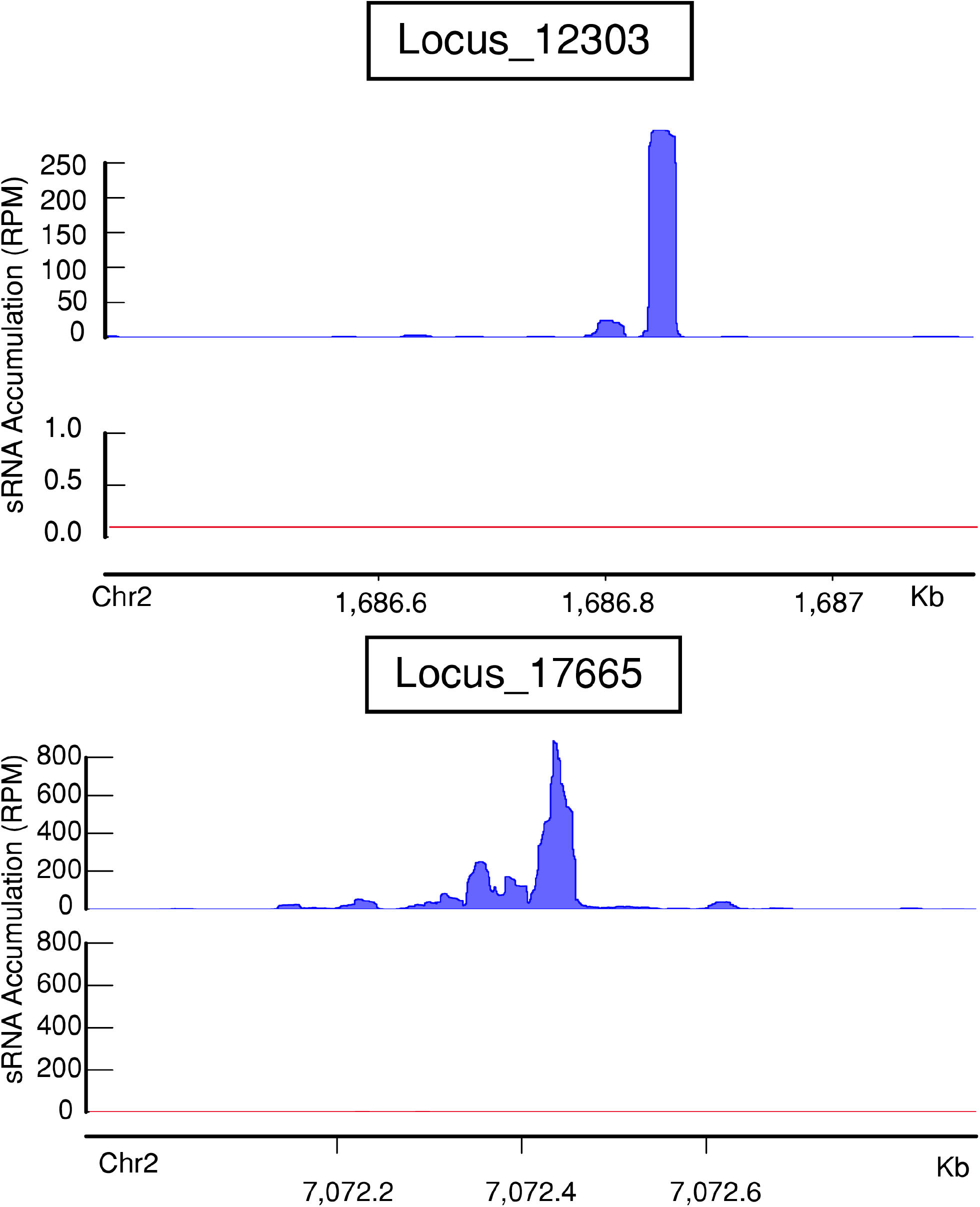
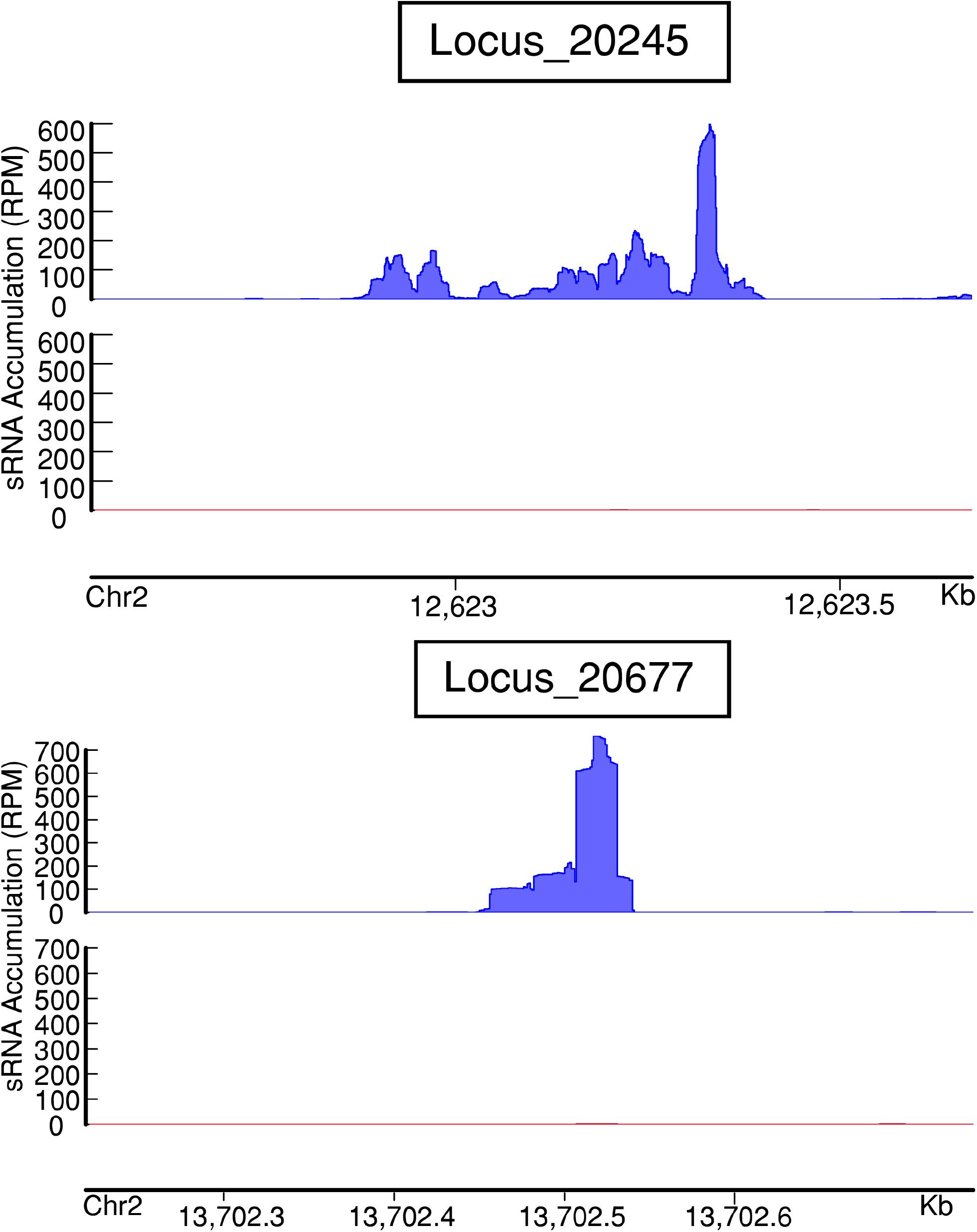
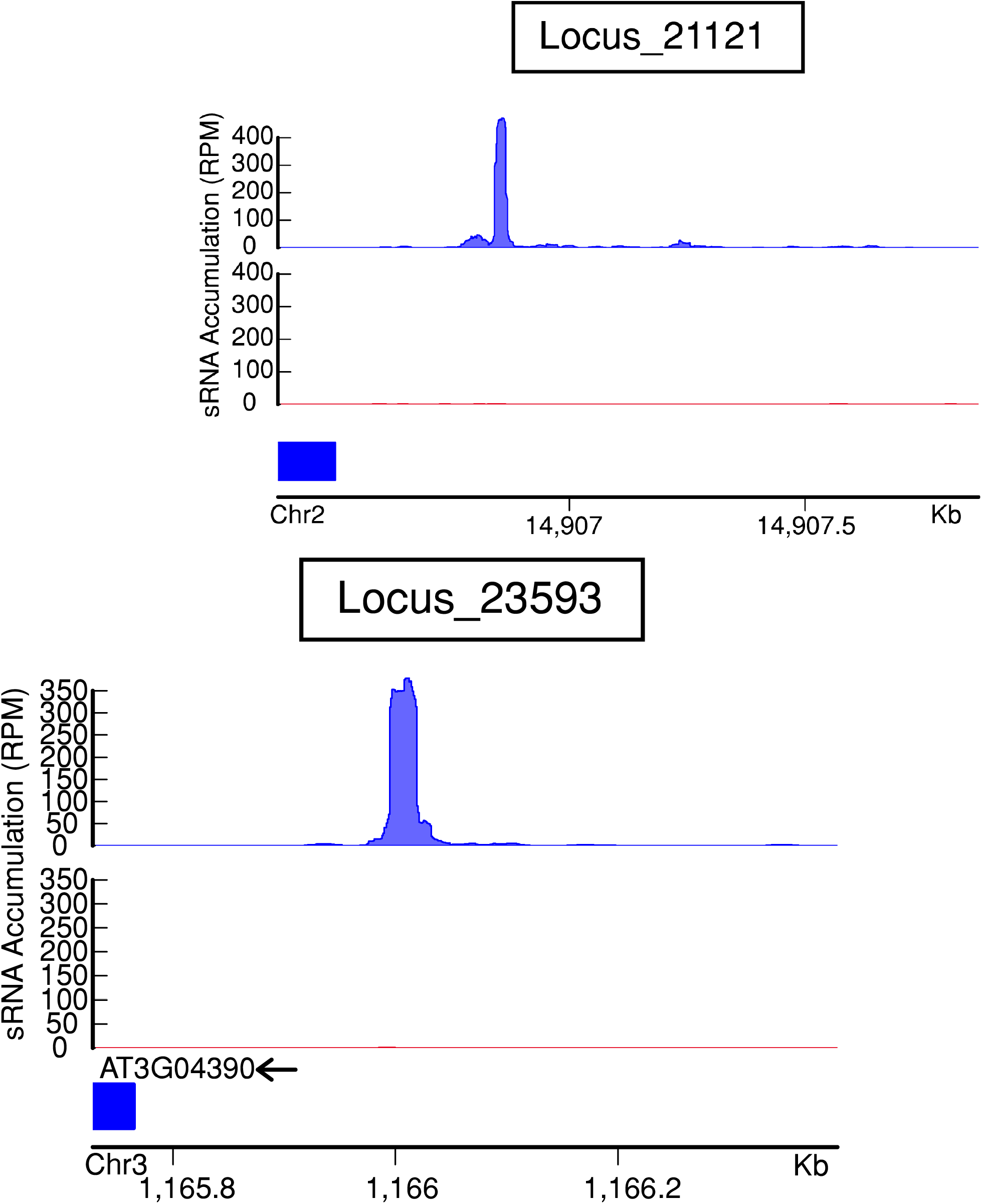
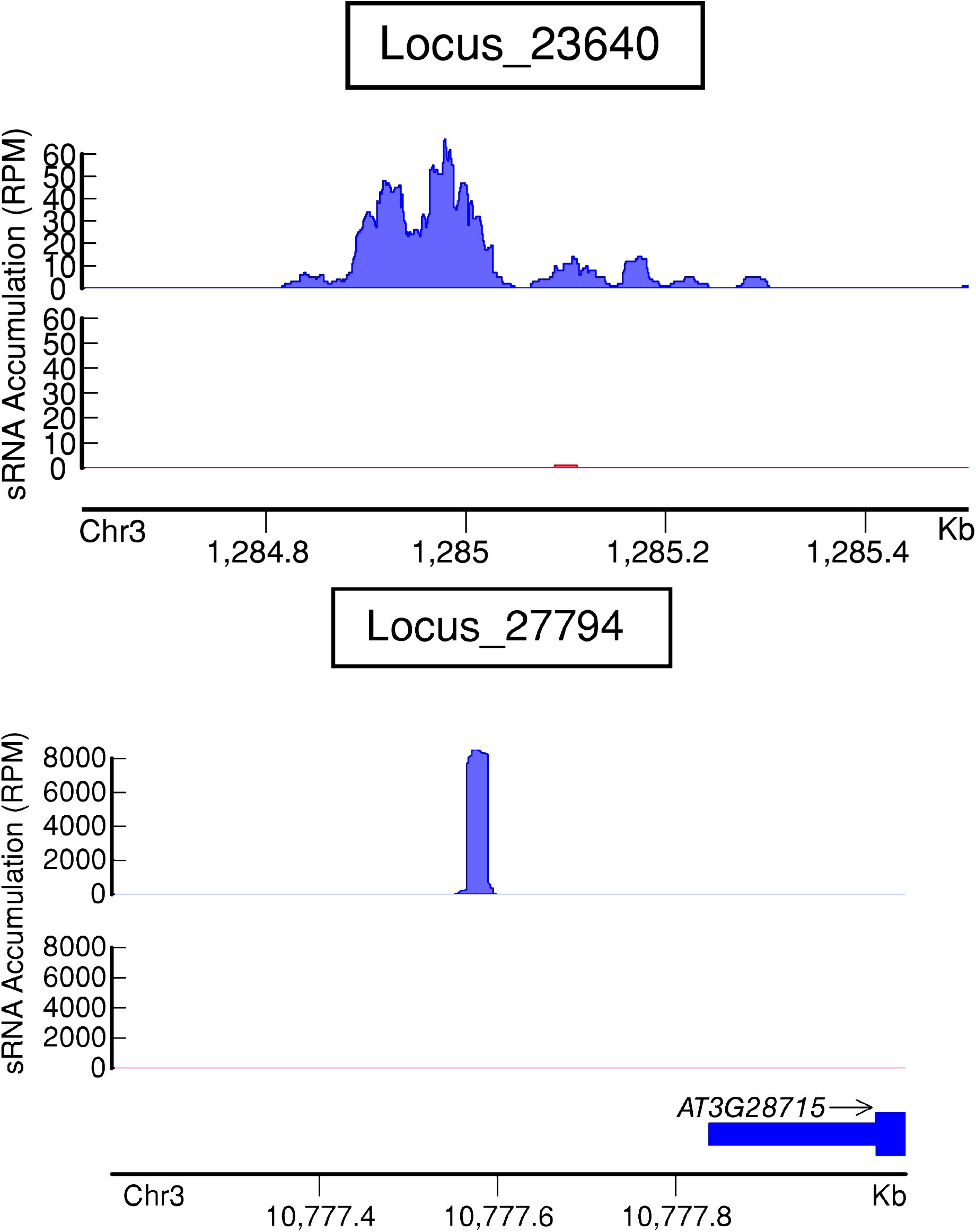
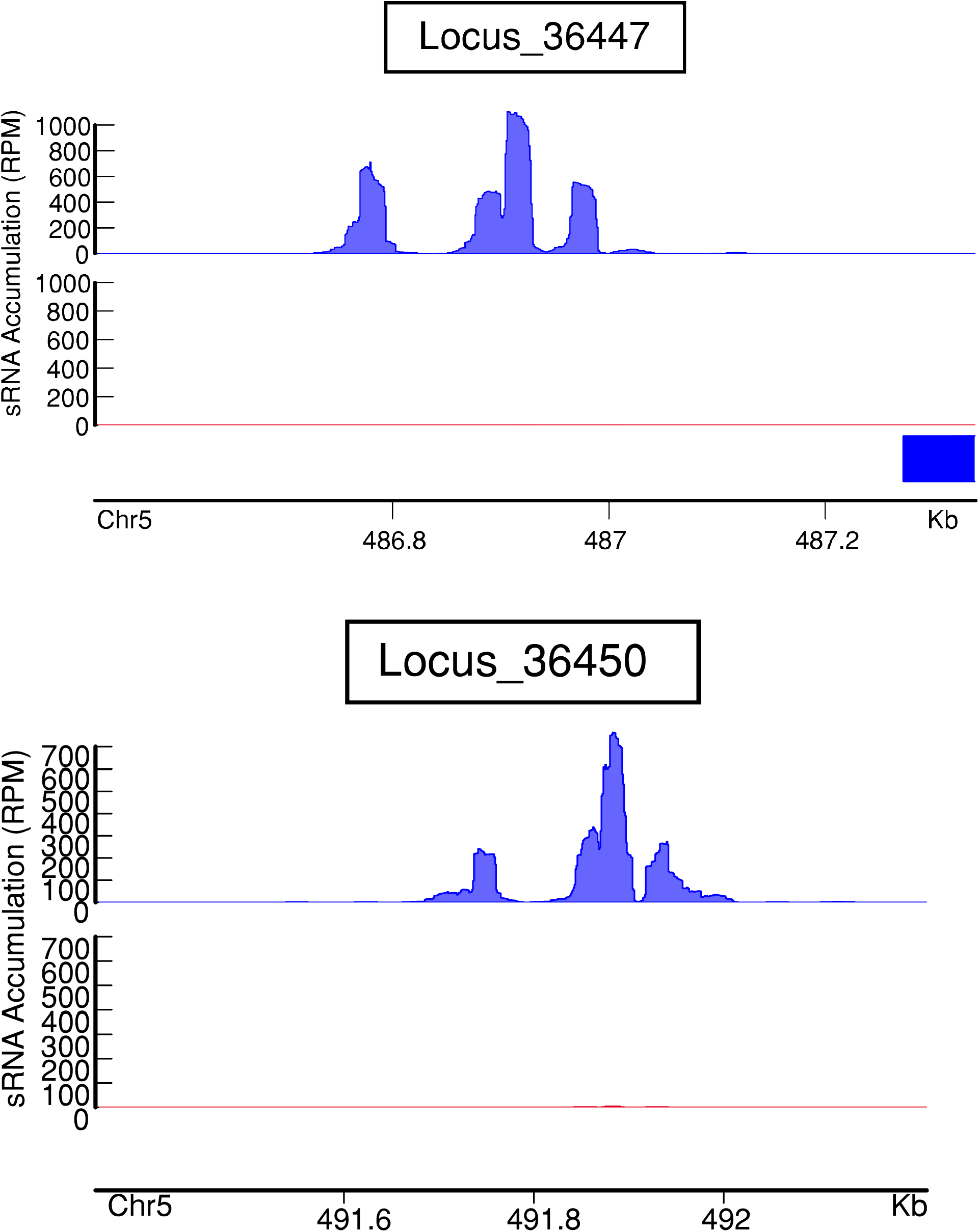
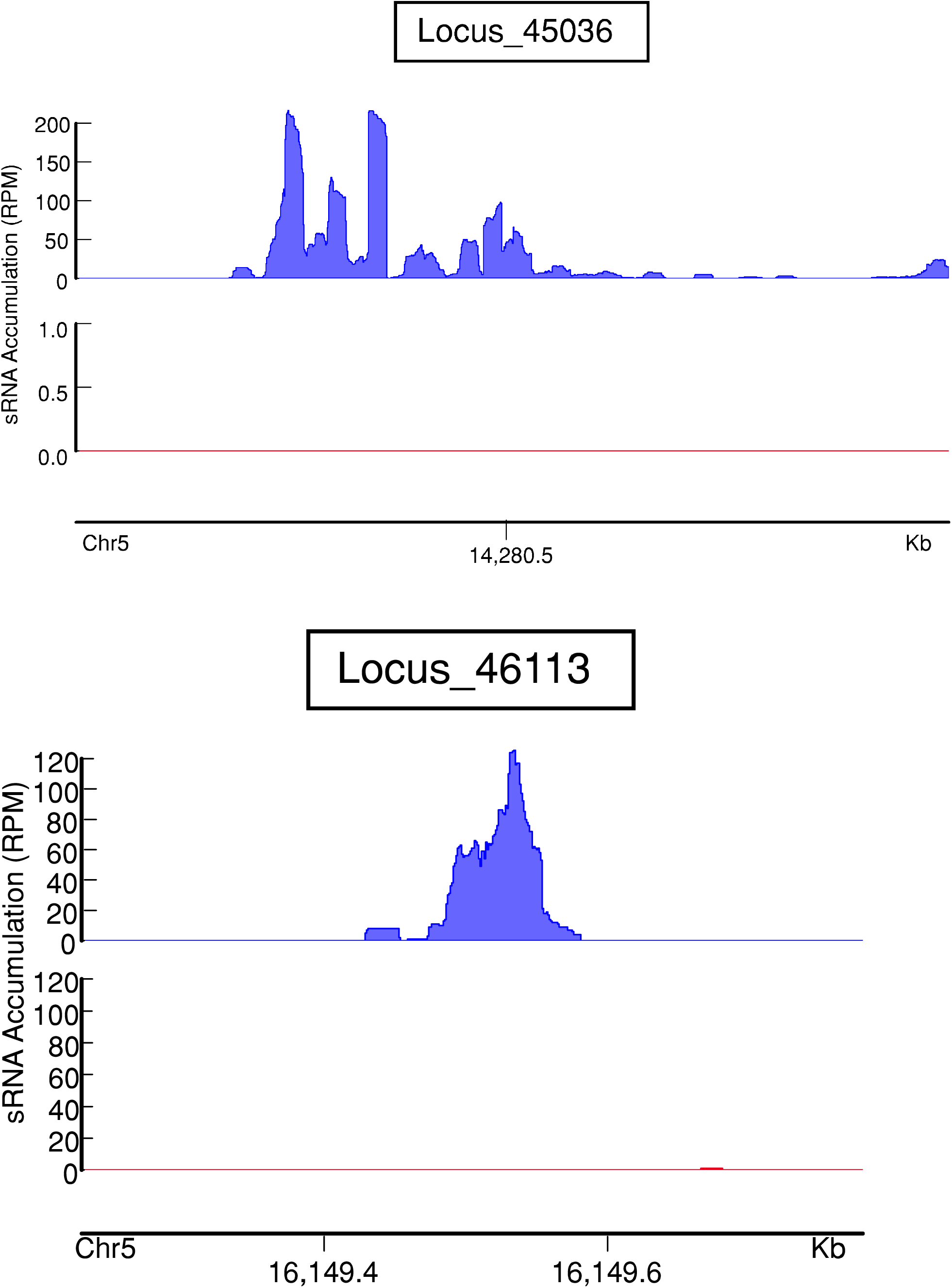
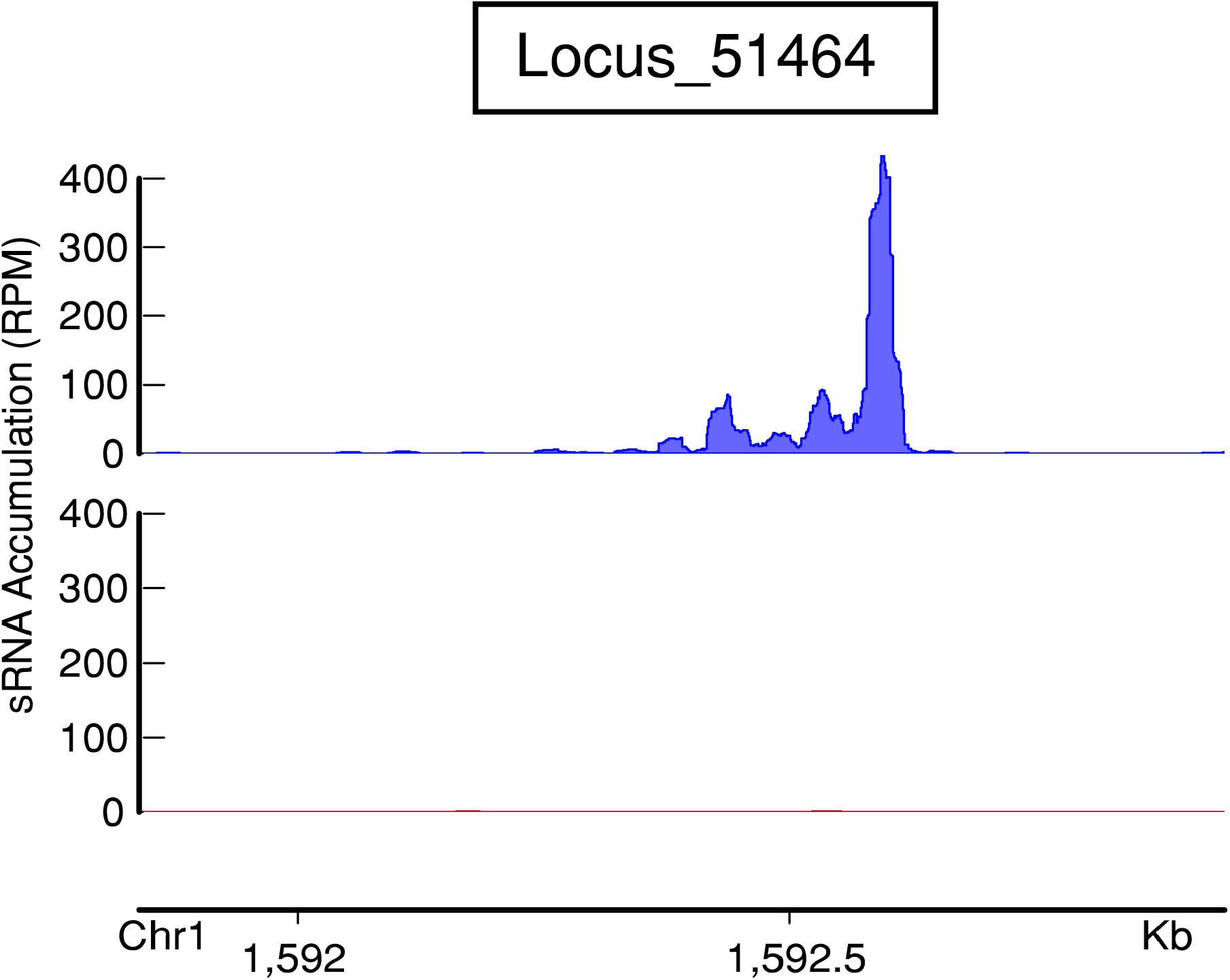
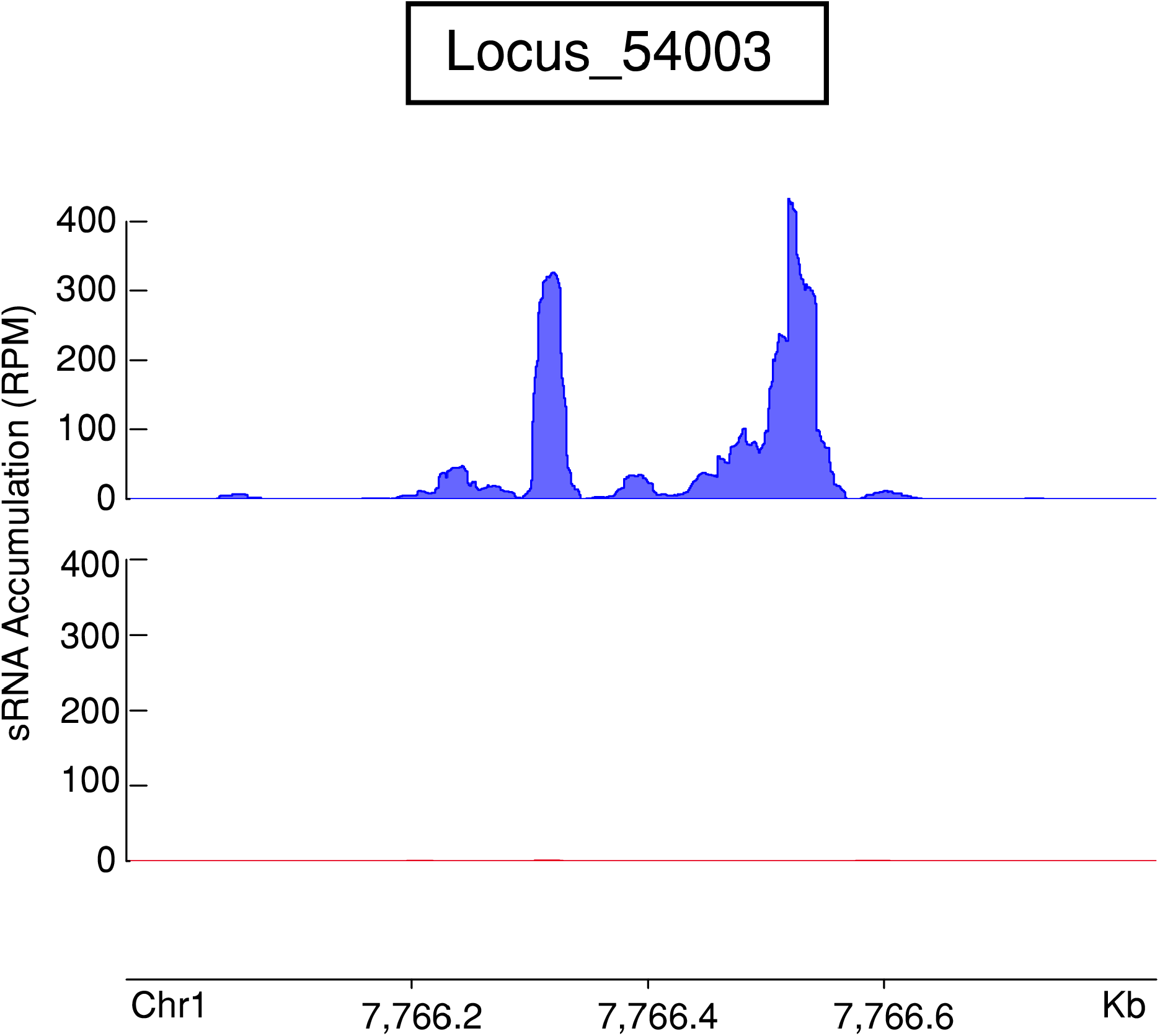
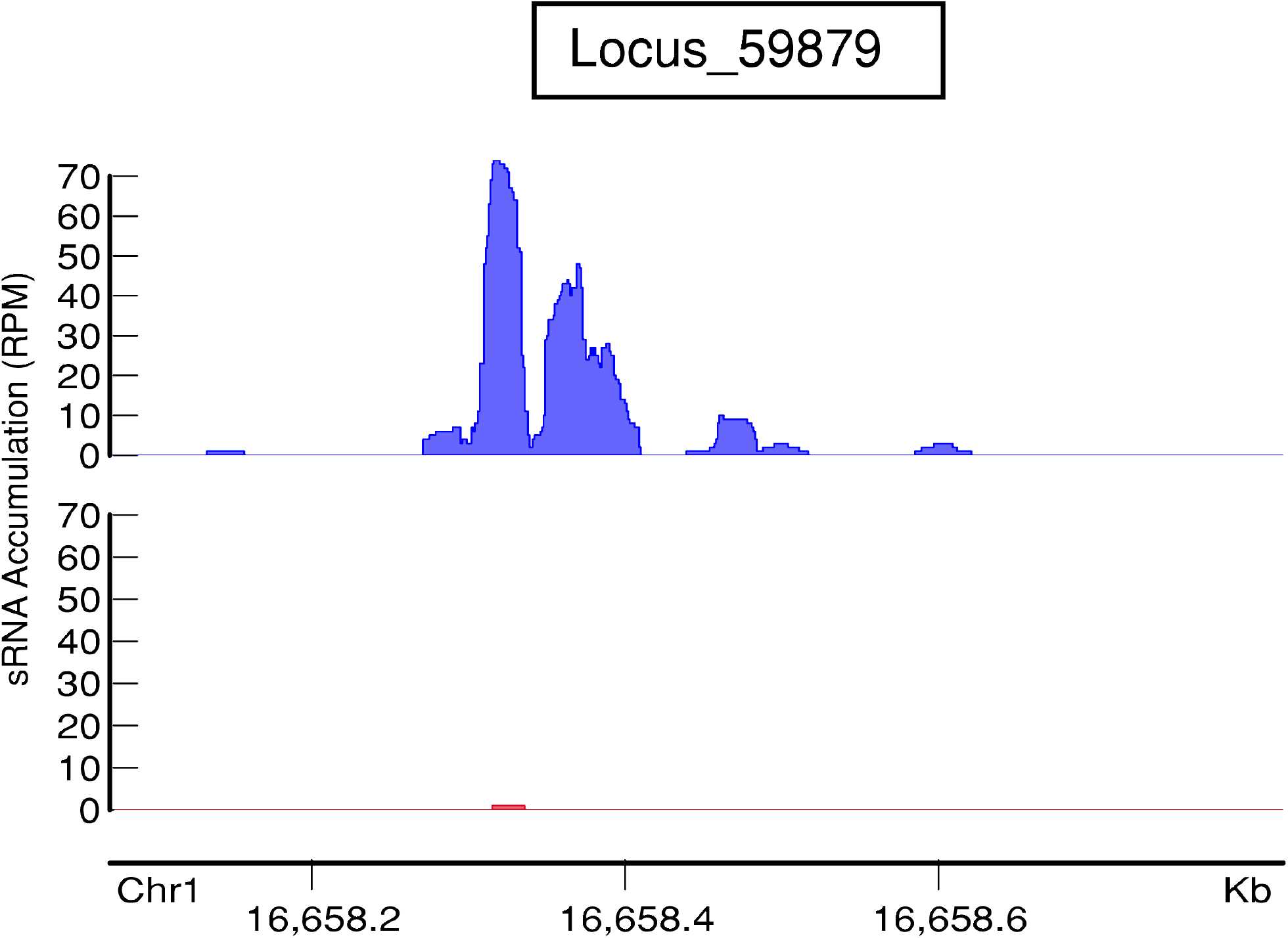
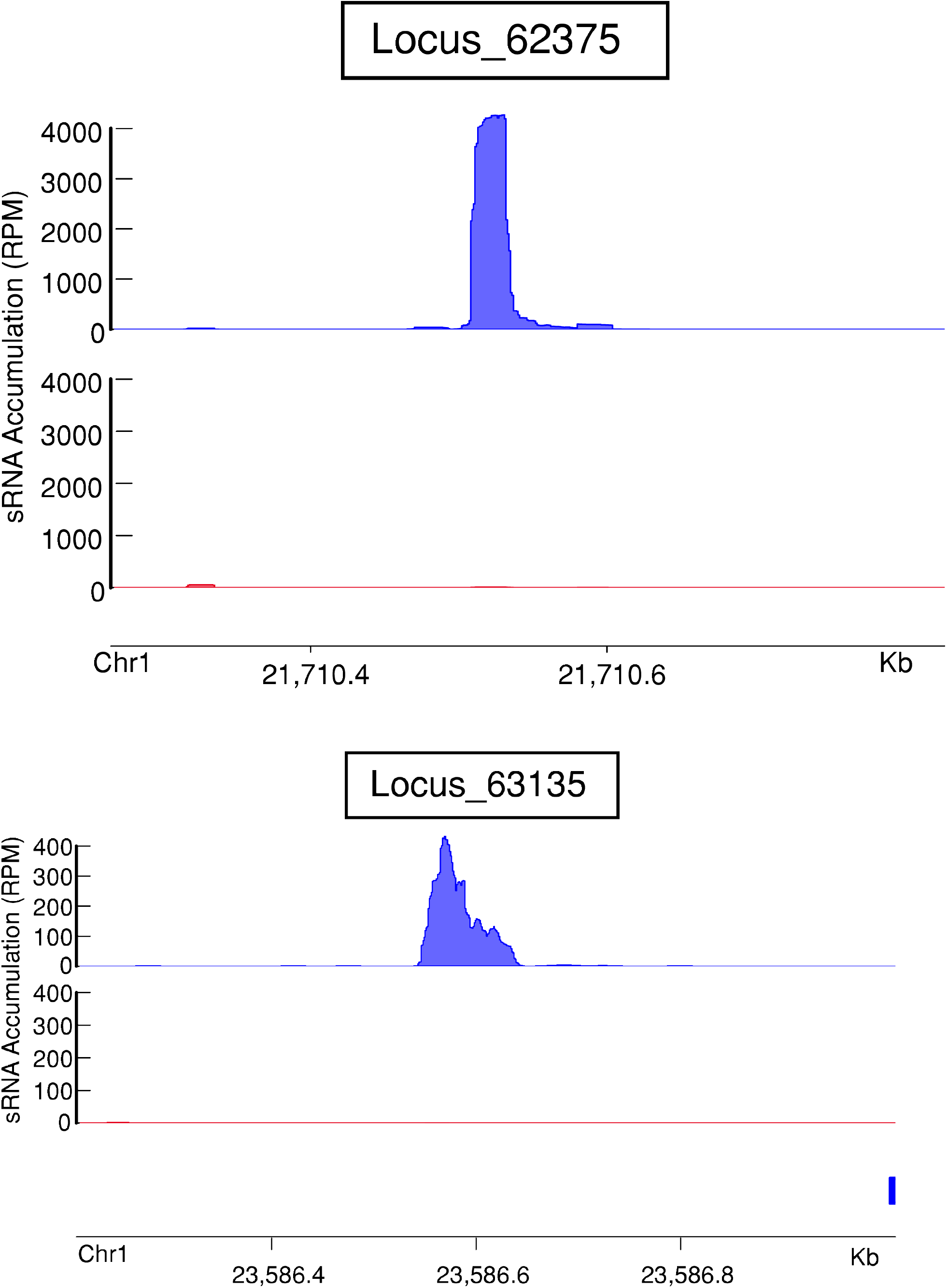
Small RNA coverage at *rdr6*-dependent, 24 nt-dominated small RNA loci. Charts represent small RNA accumulation in RPMs at a locus (regardless of the strandedness of a read). Top chart (blue) represents accumulation in wild-type libraries, and the bottom chart (red) represents the same in *rdr1/2/6* libraries. Arrows on the gene names represent the strandedness of that gene. None had significant phasing.

**Figure S9.**
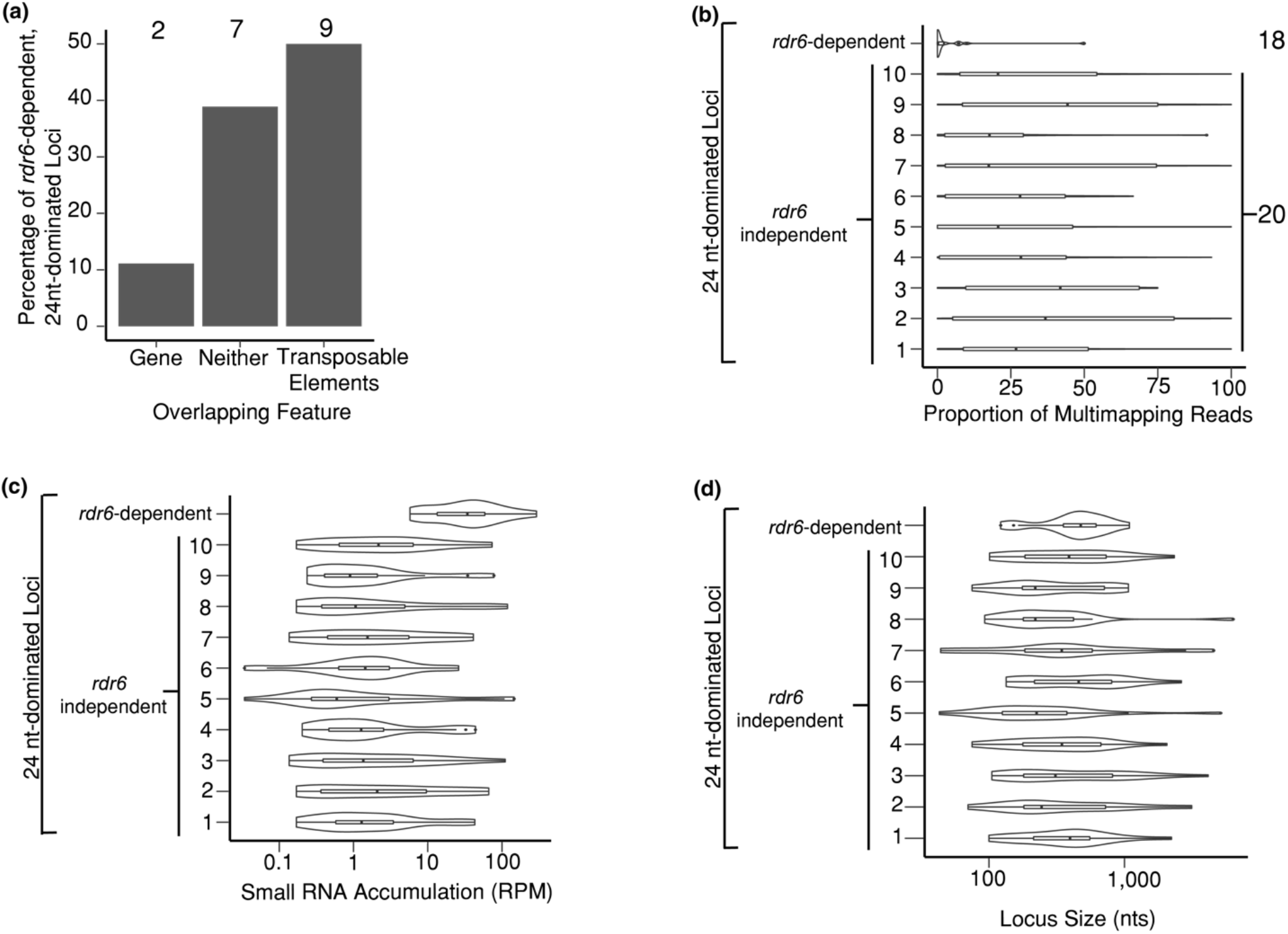
*rdr6*-dependent, 24 nt-dominated loci’ characteristics compared to other 24nt-dominated loci. (a) The overlap between *rdr6*-dependent, 24 nt-dominated loci with genes and transposons was calculated as in Figure 3a. Numbers at the top indicate the count in each category. (b) The proportion of multi-mapping reads produced at *rdr6*-dependent, 24 nt-dominated loci compared to other 24 nt-dominated loci. Numbers at the top indicate the count in each category. (c) Same as panel b except showing small RNA accumulation (in RPM). Amount in each category is the same as panel b. (d) Same as panel b except showing length (in nts). Amount in each category is the same as panel b.

**Figure S10.**
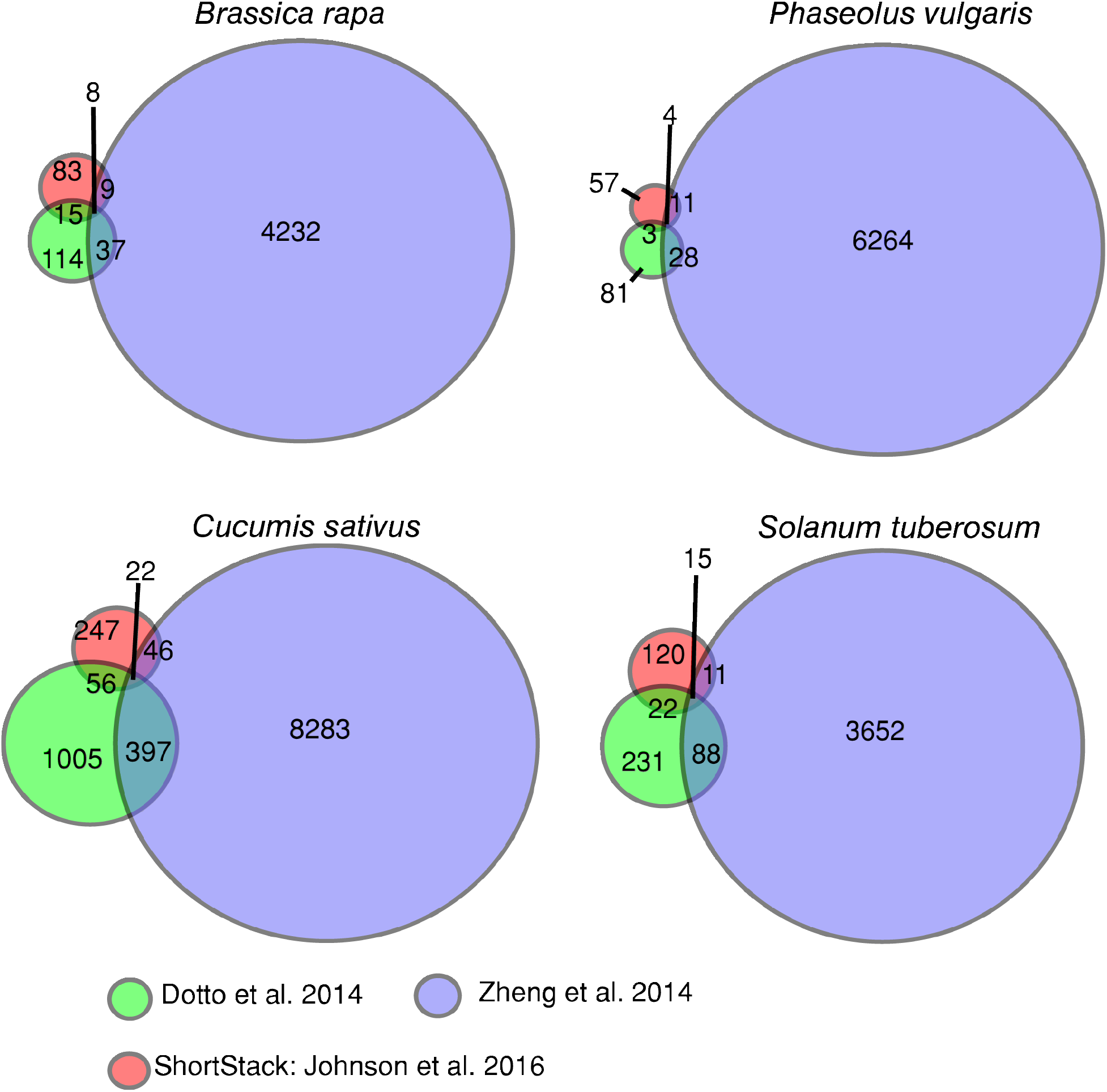
Several 24 nt-dominated small RNA loci pass *PHAS*-detection algorithms in four other eudicots. Venn diagram shows numbers of 24 nt-dominated loci that were called ‘phased’ by the indicated algorithms. Species examined is shown above the graphs.

**Table S1.**
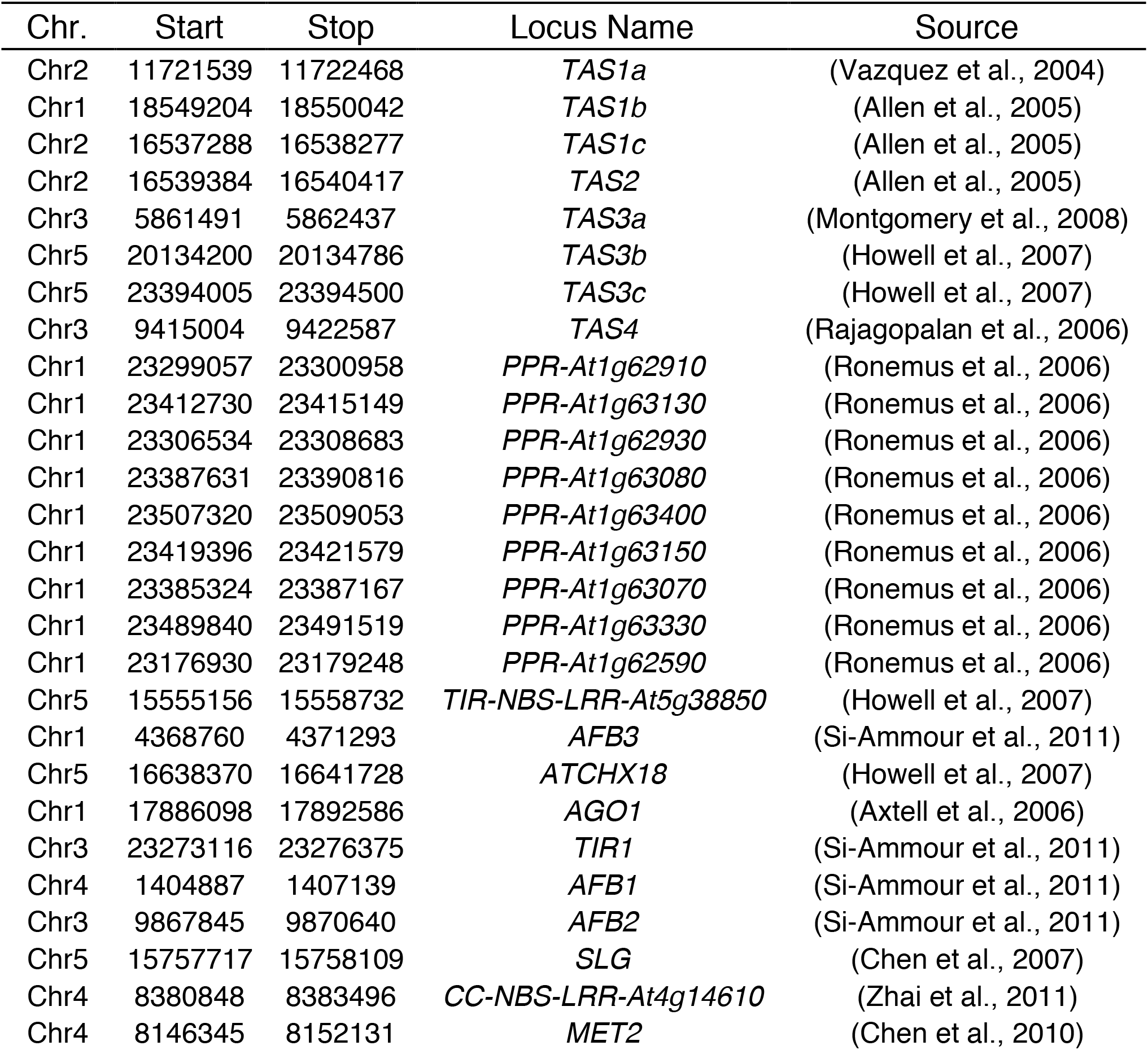
List of 25 known 21 nt *PHAS* loci in *A. thaliana*.

---

**Table S2.** *A. thaliana* small RNA libraries used in this study

| Accession Number | Genotype | 3' Adapter (First 8 nts) | Source |
| --- | --- | --- | --- |
| GSM2825283 | Wild-type Replicate 1 | TGGAATTC | (Polydore & Axtell, 2018) |
| GSM2825284 | Wild-type Replicate 2 | TGGAATTC | (Polydore & Axtell, 2018) |
| GSM2825285 | Wild-type Replicate 3 | TGGAATTC | (Polydore & Axtell, 2018) |
| GSM2825286 | *rdr1-1/2-1/6-15* Replicate 1 | TGGAATTC | (Polydore & Axtell, 2018) |
| GSM2825287 | *rdr1-1/2-1/6-15* Replicate 2 | TGGAATTC | (Polydore & Axtell, 2018) |
| GSM2825288 | *rdr1-1/2-1/6-15* Replicate 3 | TGGAATTC | (Polydore & Axtell, 2018) |
| GSM1533527 | Wild-type Replicate 1 | TGGAATTC | (Groth et al., 2014) |
| GSM1533528 | Wild-type Replicate 2 | TGGAATTC | (Groth et al., 2014) |
| GSM1533529 | Wild-type Replicate 3 | TGGAATTC | (Groth et al., 2014) |
| GSM1533542 | *dcl3* Replicate 1 | TGGAATTC | (Groth et al., 2014) |
| GSM1533543 | *dcl3* Replicate 2 | TGGAATTC | (Groth et al., 2014) |
| GSM1533544 | *dcl3* Replicate 3 | TGGAATTC | (Groth et al., 2014) |
| GSM1845210 | Wild-type Replicate 1 | AGATCGGA | (Elvira-Matelot et al., 2016) |
| GSM1845211 | Wild-type Replicate 2 | AGATCGGA | (Elvira-Matelot et al., 2016) |
| GSM1845212 | Wild-type Replicate 3 | AGATCGGA | (Elvira-Matelot et al., 2016) |
| GSM1845222 | *dcl2-1/3-1/4-2t* Replicate 1 | AGATCGGA | (Elvira-Matelot et al., 2016) |
| GSM1845223 | *dcl2-1/3-1/4-2t* Replicate 2 | AGATCGGA | (Elvira-Matelot et al., 2016) |
| GSM1845224 | *dcl2-1/3-1/4-2t* Replicate 3 | AGATCGGA | (Elvira-Matelot et al., 2016) |
| GSM1087973 | Wild-type Replicate 1 | TCGTATGC | (Jeong et al., 2013) |
| GSM1087974 | Wild-type Replicate 2 | TCGTATGC | (Jeong et al., 2013) |
| GSM1087975 | *dcl1-7* Replicate 1 | TCGTATGC | (Jeong et al., 2013) |
| GSM1087976 | *dcl1-7* Replicate 2 | TCGTATGC | (Jeong et al., 2013) |
| GSM1377370 | Wild-type Replicate 1 | TGGAATTC | (Li et al., 2014) |
| GSM1377371 | Wild-type Replicate 2 | TGGAATTC | (Li et al., 2014) |
| GSM1377372 | *nrpd1-3* Replicate 1 | TGGAATTC | (Li et al., 2014) |
| GSM1377373 | *nrpd1-3* Replicate 2 | TGGAATTC | (Li et al., 2014) |
| GSM1377376 | *rdr2-1* Replicate 1 | TGGAATTC | (Li et al., 2014) |
| GSM1377377 | *rdr2-1* Replicate 2 | TGGAATTC | (Li et al., 2014) |
| GSM2102962 | Wild-type Replicate 1 | TGGAATTC | (Panda et al., 2016) |
| GSM2102963 | Wild-type Replicate 2 | TGGAATTC | (Panda et al., 2016) |
| GSM2102965 | *rdr6-15* Replicate 1 | TGGAATTC | (Panda et al., 2016) |
| GSM2102462 | *rdr6-15* Replicate 2 | TGGAATTC | (Panda et al., 2016) |
| GSM893112 | Wild-type Replicate 1 | CACTCGGG | (Lee et al., 2012) |
| GSM893113 | Wild-type Replicate 2 | CACTCGGG | (Lee et al., 2012) |
| GSM893114 | Wild-type Replicate 3 | CACTCGGG | (Lee et al., 2012) |
| GSM893115 | *nrpb1-1 (nrpe)* Replicate 1 | CACTCGGG | (Lee et al., 2012) |
| GSM893116 | *nrpb1-1 (nrpe)* Replicate 2 | CACTCGGG | (Lee et al., 2012) |
| GSM893117 | *nrpb1-1 (nrpe)* Replicate 3 | CACTCGGG | (Lee et al., 2012) |
| GSM1668899 | Wild-type Replicate 1 | TGGAATTC | (Zhai et al., 2015) |
| GSM1668905 | Wild-type Replicate 2 | TGGAATTC | (Zhai et al., 2015) |

